# Urbanisation generates multiple trait syndromes for terrestrial taxa worldwide

**DOI:** 10.1101/2023.05.24.541105

**Authors:** Amy K. Hahs, Bertrand Fournier, Myla F. J. Aronson, Charles H. Nilon, Adriana Herrera-Montes, Allyson Salisbury, Caragh G. Threlfall, Christine C. Rega-Brodsky, Christopher A. Lepczyk, Frank A La Sorte, Ian MacGregor-Fors, J. Scott MacIvor, Kirsten Jung, Max R. Piana, Nicholas S.G. Williams, Sonja Knapp, Alan Vergnes, Aldemar A. Acevedo, Alison M. Gainsbury, Ana Rainho, Andrew J. Hamer, Assaf Shwartz, Christian C. Voigt, Daniel Lewanzik, David M. Lowenstein, David O’Brien, Desiree Tommasi, Eduardo Pineda, Ela Sita Carpenter, Elena Belskaya, Gabor Lövei, James C Makinson, Jennifer Castañeda-Oviedo, Joanna Coleman, Jon P. Sadler, Jordan Shroyer, Julie Teresa Shapiro, Katherine C. R. Baldock, Kelly Ksiazek-Mikenas, Kevin C. Matteson, Kyle Barrett, Lizette Siles, Luis F. Aguirre, Luis Orlando Armesto, Marcin Zalewski, Maria Isabel Herrera-Montes, Martin K. Obrist, Rebecca K. Tonietto, Ricardo Torrado, Sara A. Gagné, Sarah J. Hinners, Tanya Latty, Thilina D. Surasinghe, Thomas Sattler, Werner Ulrich, Tibor Magura, Zoltan Elek, D. Johan Kotze, Marco Moretti

**Author notes:** Joint First Authors (Amy K. Hahs, Bertrand Fournier). Joint Last Authors.

## Abstract

Cities can host significant biological diversity. Yet, urbanisation leads to the loss of habitats and, potentially, to local extinctions. Understanding how multiple taxa respond to urbanisation globally is essential to promote and conserve biodiversity in cities and surrounding landscapes. Using a dataset with site-level occurrence and trait data of 5302 species from six terrestrial fauna taxonomic groups across 379 cities on 6 continents, we show that urbanisation produces taxon-specific changes in trait composition, with traits related to reproductive strategy consistently showing the strongest response. The effect of urbanisation on community trait composition is strongest at the largest spatial scale considered, and more closely linked to landscape composition (% urban) than arrangement (aggregation), although latitude and climatic variables remain a stronger influence. This study did not find evidence in support of a global urban taxa syndrome, but instead we suggest that there are four general urban trait syndromes, with resources associated with reproduction and diet likely to be driving patterns in traits associated with mobility and body size. Functional diversity measures showed a wide range of responses, leading to a shift in trait space that is most likely driven by the distribution and abundance of critical resources, and the urban trait syndrome displayed by individual species within a community. Further research is required to understand the interactions between the four general urban trait syndromes, resource distribution and abundance and changes in functional diversity of taxa at different spatial and temporal scales. Maximising opportunities to support species within taxa groups with different urban trait syndromes should be pivotal in conservation and management programmes within and among cities. This will reduce the likelihood of biotic homogenisation at the taxa level, and helps ensure that urban environments have the ecological capacity to respond to challenges such as climate change, further habitat fragmentation and loss, and other disruptions. These actions are critical if we are to reframe the role of cities in global biodiversity loss.

## Introduction

Cities across the globe host significant biological diversity^1–2^ that provide key ecosystem services for over 50% of the world’s human population^3^. Urban growth often coincides with regional and global biodiversity hotspots^4^ and occurs fastest in low-elevation, biodiversity-rich coastal zones^5^. Thus, although urban environments cause significant loss and transformation of habitats and modify landscape spatial structure, minimising these impacts will be critical if we are to counter their role in the current extinction crisis^6^. Understanding how multiple taxa respond, through their functional traits, to the environmental pressures and filters of urbanisation globally is essential to formulate effective strategies to promote biodiversity in urban environments.

Although considerable progress has been made toward understanding the impacts of urbanisation on global biodiversity, certain key research gaps remain. The scientific literature is geographically biased towards larger metropolitan areas^7^ of the Northern Hemisphere and Australia^5^. Meanwhile, most biodiversity hotspots are in the tropics and the Southern Hemisphere and have received less attention^8^. Urban landscape structure has largely been characterised by negative aspects such as the proportion of impermeable surfaces, whereas the enabling aspects for biodiversity such as spatial configuration and the proportion of vegetation cover are relatively understudied^9^, especially at the global level. Urban biodiversity studies are also heavily biased taxonomically towards plants and birds^10^. Other speciose and functionally-important groups, such as insects, amphibians, bats and reptiles are severely impacted by urbanisation but poorly studied^11–14^. Despite the increasing importance of functional traits in the ecological literature and recent efforts to integrate functional aspects of biodiversity into urban ecological research^15^, most urban biodiversity investigations remain focused on taxonomic diversity^16^. This hampers our ability to develop a mechanistic understanding of the impact of urbanisation on biodiversity; creates additional challenges when making cross-taxa or cross- region comparisons^17^; and hinders our ability to effectively conserve species with different life histories and habitat requirements.

Traits are the attributes of a species that describe morphology, phenology, behaviour, and life history and influence all aspects of an organism’s fitness^18^. Trait-based approaches make it possible to characterise the functional aspects of biodiversity^19^. They facilitate cross-taxa and cross-region comparisons^20^, and provide insights into the ecological processes driving species assemblages^21^. Trait-based approaches are particularly suited to investigating the drivers of local community composition, including environmental filtering and biotic interactions^22–23^. Such knowledge is critical to the understanding and proactive management of the effects of urbanisation on biodiversity and its associated ecological functions and ecosystem services.

Cities impose strong filters on local faunal assemblages ranging from habitat loss to changes in local climate and environmental conditions and novel habitats and species interactions^24^. This filtering process is hypothesised to lead to global biotic, taxonomic and functional homogenisation, such that well-adapted species with similar traits or life histories become increasingly widespread geographically and locally abundant^25–27^. Cosmopolitan generalist species are found in most cities around the world^1^, while specialist species tend to disappear^28^. Although exceptions exist, cities tend to select for small and highly mobile fauna that have a broad environmental niche and a generalist diet^15, 29, 30^. While evidence for global functional homogenisation remains inconclusive due to different legacies and regional species pools, leading to high variability of local biodiversity in cities^31^, current understanding suggests that *highly urbanised environments favour **mobile** and **r-reproductive strategist** species with a **generalist diet**, leading to a decrease in functional diversity*. We hypothesise that increased representation of these traits across multiple taxa in cities around the world supports the proposition that there is an ‘urban syndrome’ associated with species’ responses to urbanisation^27^. This study sets out to:

1. Test our hypothesis by evaluating evidence against the current understanding of an ‘urban syndrome’ related to average community traits and/or functional diversity;
2. Investigate whether the proportion and spatial aggregation of urban land and forest cover (see Methods) induce stronger changes in community functional diversity than known latitudinal or climatic trends. In this case, we use urban land cover to represent a gradient of urbanisation filters, and forest cover to represent the amount of tree canopy cover;
3. Investigate the spatial scale at which the proportion of urban land has the strongest effect, and how this differs among functional groups.

This study used a collaboratively compiled dataset of 5302 species found in > 70000 plots across 379 cities from 48 countries (Fig. 1) to investigate how urbanisation shapes the community trait-composition and diversity of six terrestrial animal taxonomic groups (amphibians, bats, bees, birds, carabid (ground) beetles, and reptiles) across the globe. The data are a collation of empirical studies at the highly-resolved spatial scale of individual sites rather than generalised to city. Only one taxa (birds) was extracted from a global biodiversity dataset (eBird). We acknowledge there are still geographic biases in the data which reflect legacies of studies published prior to 2017^10^. We are also aware that there are additional taxa groups that we would have liked to include but lacked the capacity to consider in this project. However, to our knowledge, this is the most comprehensive compilation to date of urban biodiversity data for several terrestrial animal taxa at the site scale. The six taxa represent a broad range of natural histories, ecologies and behaviours and have sufficient occurrence data and trait information to conduct a global study, despite some geographic biases. The traits we considered were body size, diet, mobility and reproductive strategy, as these are all important for an individuals’ survival, growth and reproduction^18^. Functional diversity metrics captured key facets of trait diversity (functional richness – FRic, functional evenness – FEve, functional dispersion - FDis), to investigate whether there was evidence to support a contraction of trait space associated with the urban syndrome. Further details can be found in the Methods and Supplementary Materials.

**Figure 1.**
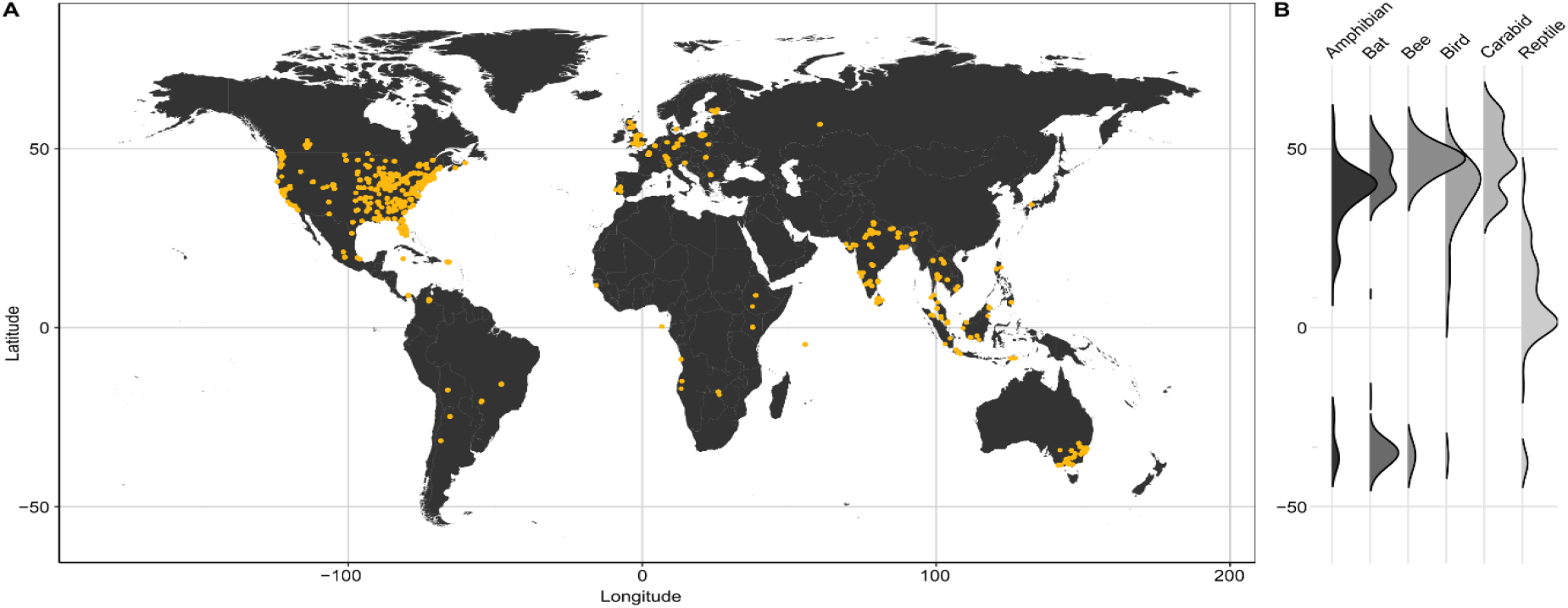
Global distribution of data included in this study. (A) Locations of sampling plots for all six taxonomic groups combined. All data are from the UrBioNet contributor network except for birds (eBird). (B) Ridgeline plots showing the density of sampling locations per taxon as a function of latitude. See Supplementary Figure S1 for taxa specific maps.

## Results

Our global analysis shows that urbanisation is a major driver of urban community functional composition. All traits and functional diversity metrics changed with increasing urban land cover, although the strength and direction of change within each trait category differed among taxa (Fig. 2).

**Figure 2.**
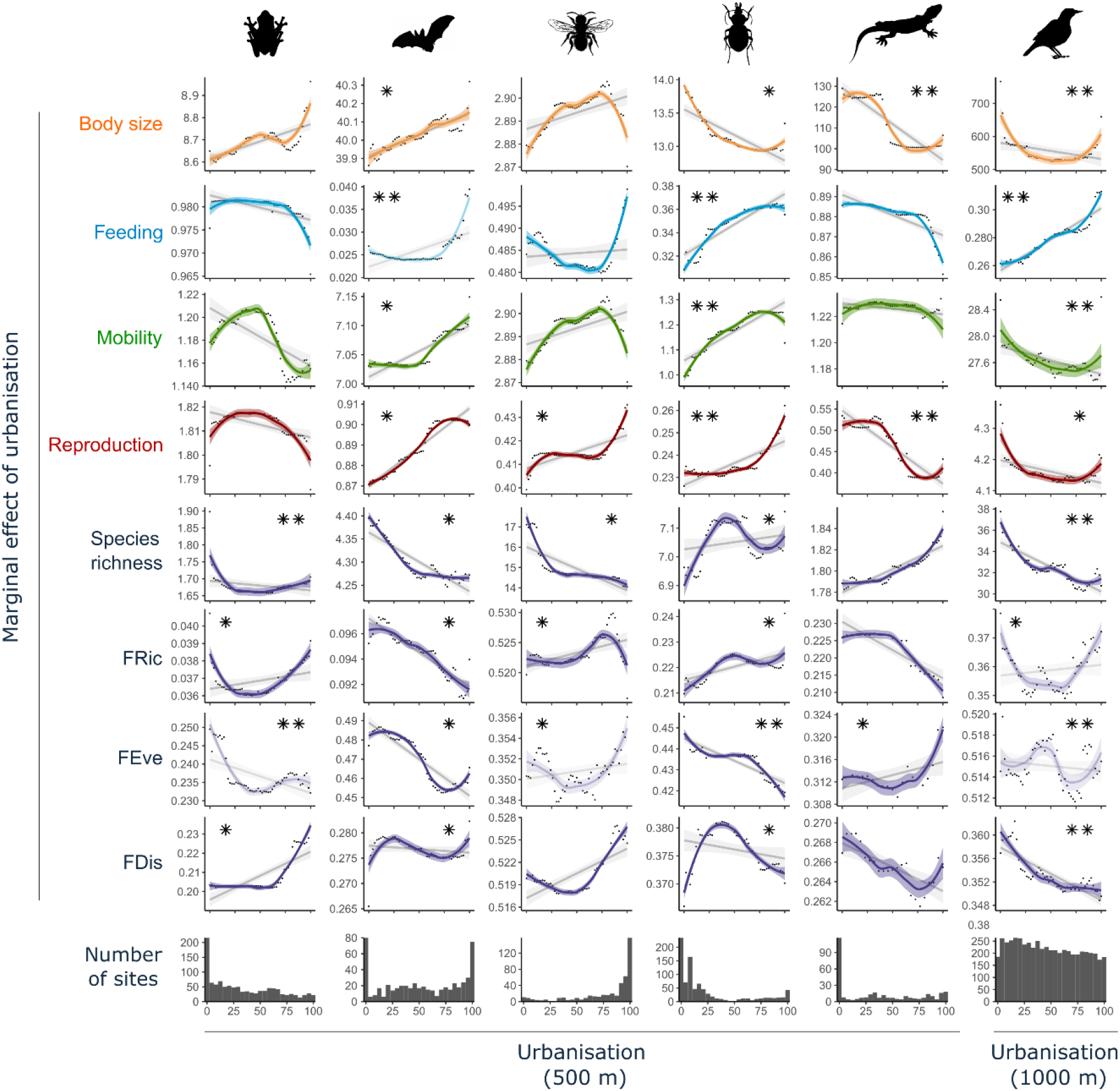
Predicted changes in trait values per taxon along an urbanisation gradient. Partial dependence plots showing the urbanisation-induced shifts in community functional metrics for six taxonomic groups. The partial dependence plots summarise the marginal effect that urbanisation (x-axis = percentage of urbanised area in a 500 m radius around the sampling plot; or 1000 m for birds) has on the predicted values of each community-level trait (i.e. effect of urbanisation when climate, latitude and forest cover are kept constant). The y-axes reflect the range of predicted values for each response variable and are not zeroed so care should be taken when interpreting the magnitude of change. The fitted colour lines and 95% confidence bands are from Local Polynomial Regression (LOESS). The grey lines are from linear regressions based on the same data to indicate direction of trend. Trait definitions are provided in Supplementary Table S3 (briefly, *Feeding*: high values = generalist diet except for bats where feeding represents different hunting strategy; *Mobility*: high values = higher mobility; *Reproduction*: amphibians, birds and reptile = clutch size **/** other taxa = reproduction strategy). Note that for bees, the inter-tegula distance was used for body size and mobility, and therefore the model presented is the same for both traits. Functional dispersion (FDis), functional richness (FRic) and functional evenness (FEve) are defined in the method section in “Faunal community functional compositions” (see also Supplementary Figure S2). Transparent shade represents models with <10% variance explained. Stars show the contribution of urbanisation to the overall model (* 20-50%; ** > 50%). Additional information on each models’ overall predictive power and the contribution of the percentage of urban land cover can be found in Table 1.

**Table 1.** Performance of models predicting traits and diversity metrics. Summary statistics of random forests models of community-weighted means of traits and functional diversity metrics. *“% explained”* is the performance of the model where high values indicate that the response variable is well-predicted by urban and forest land cover, climate, and latitude. *“% explained”* was calculated as R-squared of the relationship between the predicted and the observed values of the independent test dataset. *“% inc MSE”* is the average increase in squared residuals when the variable is permuted. It represents the specific contribution (or importance) of the percentage of urban land cover (within a 500 m radius for all other taxa except birds for which we used a 1000 m radius) to the overall model performance. High values suggest that urban land cover is an important predictor.

|  |  | Amphibians | Bats | Bees | Birds | Carabids | Reptiles |
| --- | --- | --- | --- | --- | --- | --- | --- |
| Body size | % explained | 62 | 44 | 32 | 18 | 40 | 62 |
|  | % inc MSE | 17 | 25 | 12 | 62 | 47 | 62 |
| Feeding | % explained | 67 | 9 | 55 | 20 | 19 | 55 |
|  | % inc MSE | 16 | 52 | 13 | 57 | 83 | 13 |
| Mobility | % explained | 17 | 31 | 32 | 16 | 46 | 42 |
|  | % inc MSE | 14 | 42 | 12 | 62 | 71 | 8 |
| Reproduction | % explained | 62 | 65 | 57 | 52 | 48 | 33 |
|  | % inc MSE | 19 | 40 | 38 | 44 | 57 | 93 |
| Sp. Richness | % explained | 68 | 56 | 80 | 46 | 61 | 70 |
|  | % inc MSE | 53 | 29 | 48 | 70 | 39 | 15 |
| FDis | % explained | 53 | 54 | 26 | 16 | 18 | 29 |
|  | % inc MSE | 31 | 34 | 13 | 50 | 49 | 15 |
| FRic | % explained | 53 | 29 | 46 | 20 | 59 | 59 |
|  | % inc MSE | 39 | 23 | 36 | 38 | 40 | 18 |
| FEve | % explained | 5 | 50 | 10 | 8 | 17 | 11 |
|  | % inc MSE | 60 | 48 | 30 | 64 | 71 | 22 |

Body size and mobility were affected differently by urbanisation depending on the taxa (Fig. 2). With increasing urban land cover, carabids, birds and reptiles displayed a tendency towards species with smaller body size (7%, 23% and 27% decrease, respectively) in the most urbanised areas relative to the least urbanised areas. Carabid beetles displayed a tendency towards increased mobility (19%), while reptiles and birds tended towards reduced mobility (1%, 5%). Amphibians and bats displayed a tendency towards larger body sizes (4%, 1%) with increasing urbanisation. Amphibians displayed a 4% drop in mobility, while bats tended towards slightly higher mobility (1%) (Fig. 2). For bees, inter-tegula distance is the trait most frequently used to represent body size and mobility, and showed an inverted u-shape, where the linear trend showed a slight increase (<1%).

Our results suggest that increased urban land cover can induce a shift toward a more specialist or generalist diet depending on the taxa considered (Fig. 2). Specifically, omnivory was favoured with increasing urban land cover for birds (19%) and carabid beetles (14%). Bees showed a u-shaped response with a linear trend towards a 3% increase in the proportion of short-tongued species (Fig. 2). However, amphibians and reptiles both demonstrated shifts towards increased dietary specialisation with increasing urbanisation (8% and 5% respectively).

Reproductive traits were the first (bats and carabids) or second (amphibians, bees, birds and reptiles) most affected trait when considered across all traits for a taxon (Fig. 3). The reproductive strategy trait had the highest proportion of variance explained for four taxa, explaining 48 – 65% of the variance for bats, bees, carabids and reptiles (% explained in Table 1). The exceptions were amphibians and birds where feeding or body size (respectively) were more important. Trends indicated that increasing urban land cover was associated with reduced clutch size (amphibians, birds and reptiles), more generalist roosting (bats), overwintering (adult (imago) in carabids) and solitary nesting (bees) (Fig. 2). Bats with generalist roosting requirements increased by 3%, bees that were solitary nesters increased by 9% compared to social nesters, and carabids showed a 4% increase in the proportion of species that overwinter as adults.

**Figure 3.**
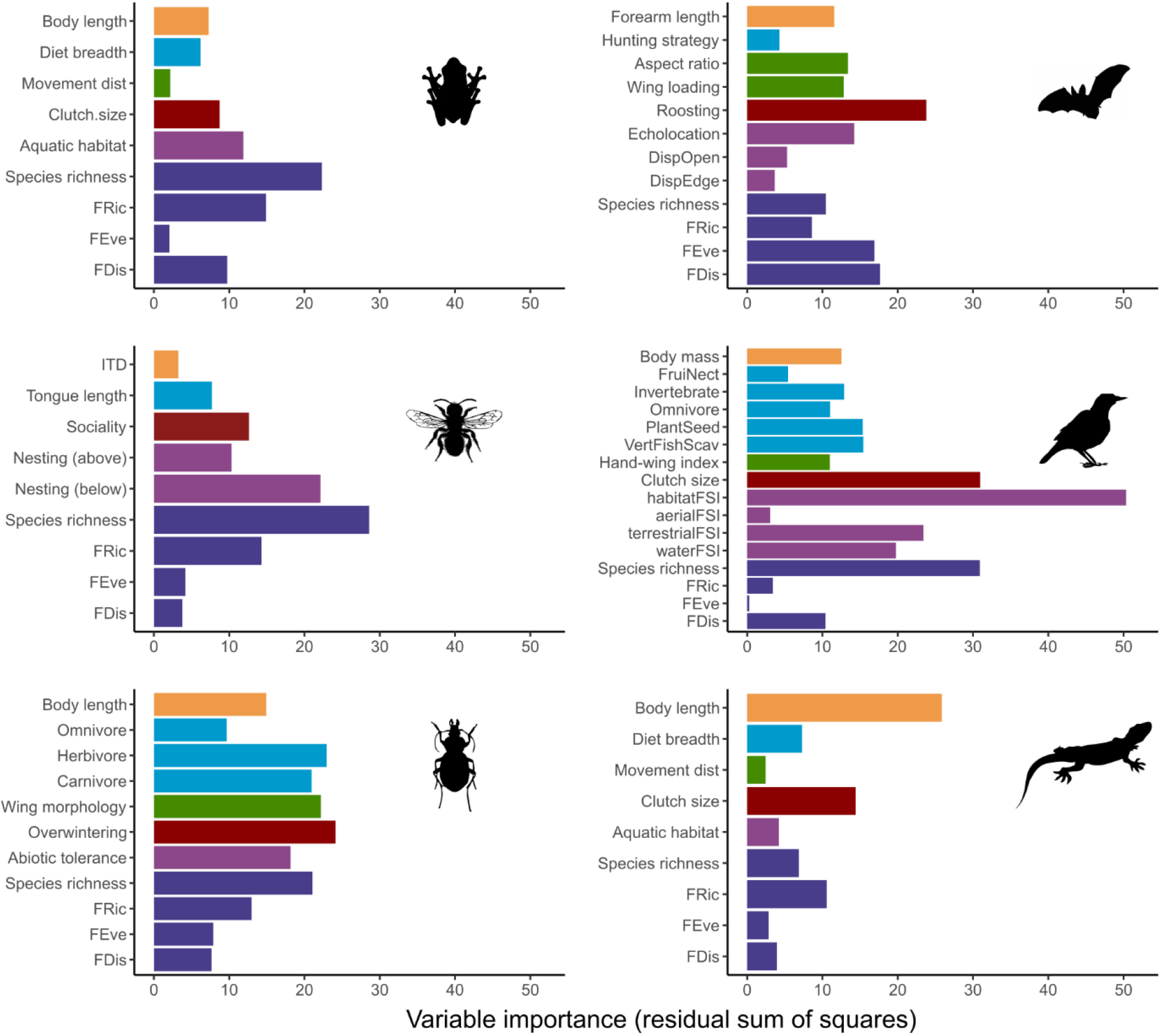
Relative importance of the extent and aggregation of urban land cover as predictors of community means (colours show the different trait categories; Supplementary Table S3) and variability (FDis = functional dispersion, FRic = functional richness, and FEve = functional evenness; dark blue) of traits as well as species richness for each taxonomic group. Variable importance was estimated using the residual sum of squares from random forests models. Average variable importance values weighted by the R^2^ of the test set of each individual model were computed to estimate urban land cover variable importance for each metric of community- weighted means and variability of traits. Longer bars indicate traits or functional diversity measures that are better predicted by urban land cover within the surrounding landscape.

The effect of urban land cover was most important at the largest spatial scale considered for all taxa examined (1000 m for birds, 500 m for all other taxa; Fig. 4). The importance of the proportion and spatial aggregation of urban land cover as predictors of taxon-specific trait syndromes ranged from 3% to 20% depending on the taxa (light blue bars, Fig. 4), but composition (%) was consistently stronger than arrangement (agg). Metrics related to forest cover were generally the least important across all taxa, with birds being the exception. Latitude and climatic region predicted shifts in community functional composition of most taxa better than urban or forest land cover or configuration. The only exception to this was again for birds, for which the importance of latitude was equal to the importance of forest cover (%) within 1000 m of the site.

**Figure 4.**
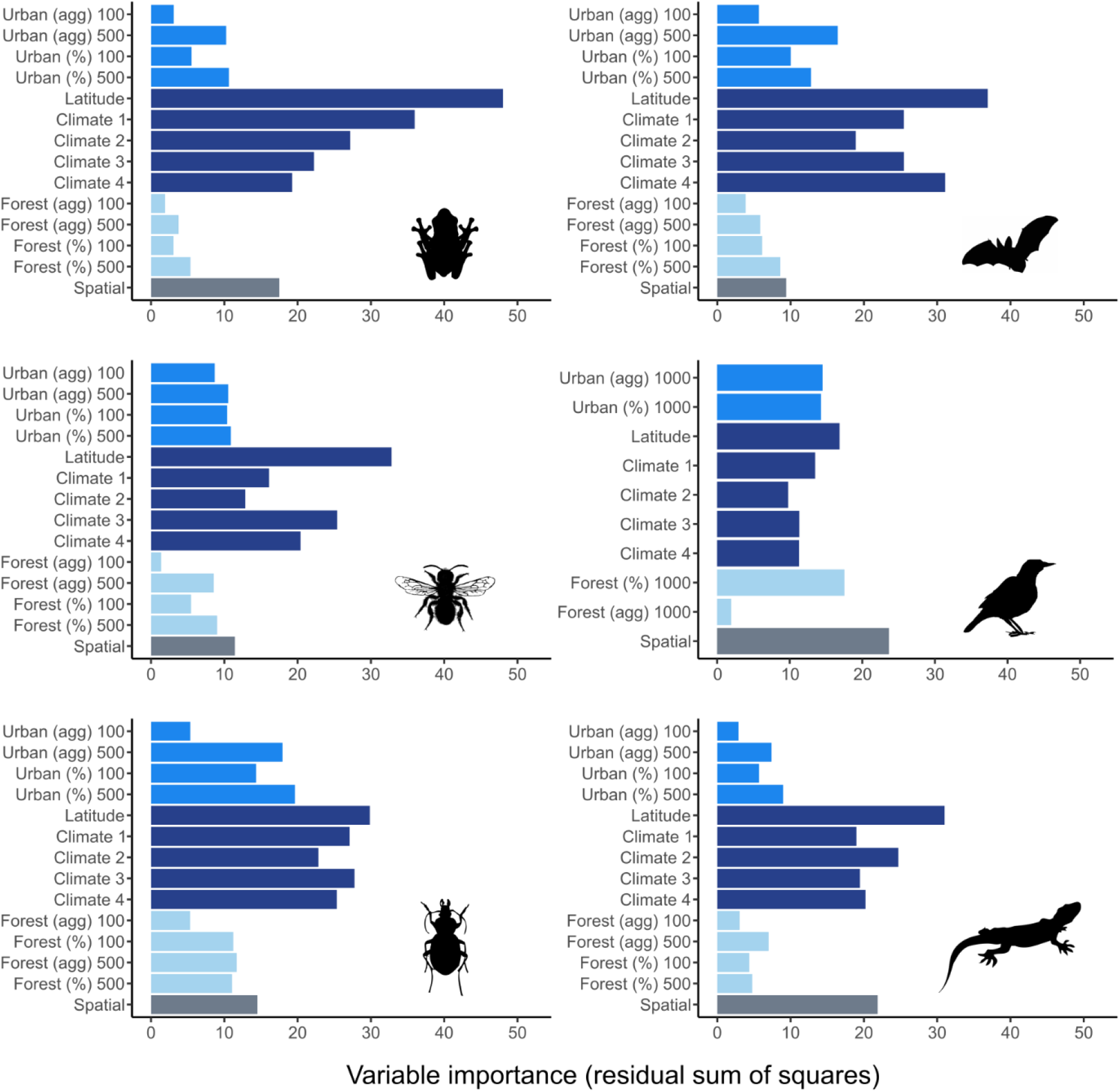
Relative importance of variables in predicting trait responses per taxon. Importance of percent cover (%) and spatial aggregation (agg) of urban and forest land cover at different buffer distances (100 m and 500 m for most taxa; 1000 m for birds), latitude, climate PCA axes, and spatial covariates (dbMEM) as predictors of the trait syndrome (i.e. considering all community weighted means and functional diversity metrics) for each taxonomic group. Variable importance was estimated using the residual sum of squares from random forests models. Average values weighted by the R^2^ of the test set of each individual model were computed to estimate variable importance for the overall trait syndromes.

There were clear effects of urbanisation on all facets of functional diversity and species richness, however they varied between taxa. Functional richness (FRic) was the functional diversity facet that was best predicted by the extent and aggregation of urban land cover for amphibians, bees, carabids and reptiles (Fig. 3; % explained in Table 1) but the direction of the response varied (Fig. 2). With increasing urbanisation, functional richness (FRic) decreased in bats (6%) and reptiles (9%), showed a u-shaped response in amphibians and birds and tended to increase in bees (2%) and carabids (8%) (Fig. 2). Functional dispersion (FDis) was a more important dimension of functional diversity for bats and birds (Fig. 3, % explained in Table 1), which declined by 4% (bats) to 5% (birds). Functional evenness (FEve), although overall poorly predicted by our models, was the dimension of functional diversity that most consistently responded strongly to urbanisation (% MSE in Table 1). Like functional diversity dimensions, species richness showed different trends depending on the taxa considered. Increasing urbanisation led to an increase in species richness of carabid beetles and reptiles (1%, 2%) but decreased the species richness all other taxa (3% amphibians, 8% bats, 18% bees, 17% birds; Fig. 2).

## Discussion

Body size and mobility are frequently correlated in functional trait studies: larger species tend to be more mobile^32^. Mobility is likely to be favoured when it helps an organism acquire resources and/or avoid competition and predation. However, our results show that for some terrestrial animal taxa, urbanisation may select for species with small home ranges that can exploit local resources^33^ and avoid risks associated with the urban matrix^34^. Reduced mobility in these taxa make them particularly vulnerable to habitat loss or degradation and can lead to the isolation of populations, increasing the importance of genetic drift and local population extinction risks.

Increasing omnivory with increasing urban land cover was observed for birds (19%) and carabid beetles (14%), which aligns with a common finding that dietary breadth predicts success in urban environments^35–36^, and our hypothesis for an ‘urban syndrome’. Bees showed a u-shaped response, which may reflect a wider diversity of flowering plants being available in urban areas, thereby providing a variety of resources for both generalist and specialist feeders. Amphibians and reptiles showed shifts towards increasing dietary specialisation. This specialisation may enable finer niche partitioning in spatially constrained spaces and thereby avoid some of the impacts of urban environments through more efficient foraging^37^. Overall, our results highlight that both generalist and specialist feeding strategies can be selected for in urban environments, but will depend on the interplay between the composition and distribution of food resources and the species ability to access and utilize them.

Our results provide evidence that urbanisation strongly selects for species with the capacity to find suitable conditions for reproduction. Fewer suitable nesting sites and higher risk of disturbance/predation in cities can thus have a strong impact on community functional composition. Providing supplemental nesting resources to compensate for loss of natural nesting possibilities can limit this impact, as has been demonstrated by the use of nest boxes to supplement the loss of hollows^38^. Increased urbanisation also influenced community mean clutch size. For example, reptiles clutch size declined by 27%, while birds displayed u-shape negative trend with 7% variation in clutch size (Fig. 2). A previous global analysis found that reptiles tend to have larger clutch sizes at higher latitudes where suitable conditions for breeding are constrained by short growing seasons or other limitations that select for reproductive strategies that maximise the number of offspring produced when food availability peaks^39^. In cities, the reduction in frost days due to the urban heat island and the greater consistency of food and water throughout the year due to horticultural plantings and human activities, may benefit species that have multiple but smaller clutches to avoid population density pressures on locally limited resources. Smaller clutch sizes in urban birds have been associated with higher survival and increased growth^40^. Reduced clutch sizes in birds have also been linked to perceptions of increased predation risk in altricial species where the young are fed and protected by parents when they are first born^41^. Future research could look more closely to understand to what extent the change in clutch size represents a change in the number of species exhibiting a given development type as altricial birds have smaller clutch sizes than precocial birds that require little parental care^42^.

Our results confirm the effect of latitude and climate as key drivers on the functional biodiversity of taxa observed in cities. Landcover effects were strongest at the largest spatial scales considered (1000 m for birds, 500 m for all other taxa), and the composition of the landscape (% cover) was more important than configuration (agg). These results highlight the importance of landscape-level management of urban biodiversity and the role of spatial context. They also provide additional support for our proposed general urban trait syndromes, which are highly influenced by the distribution and abundance of resources within the landscape.

We acknowledge that processes occurring at larger spatial scales than those considered in this study can also be important, especially for species with high mobility. Equally, there may also be finer scale processes that we were not able to consider due to the resolution of available datasets. Future research could address these limitations or could expand our approach to look at a wider range of taxa. The study could also be repeated in the future when empirical data from a wider range of geographic regions are available to test how well the patterns observed here continue to apply.

### Four general urban trait syndromes, rather than one universal syndrome

Our study indicates that rather than a single urban syndrome, there is strong evidence to support that each taxon has an individual urban trait syndrome each of which can be classified into one of three typologies: **mobile generalists, site specialists,** and **central place foragers** (Fig. 5), or hypothetically into a fourth typology: **mobile specialists**. The urban trait syndrome for mobile generalists most closely matches our original hypothesis that urbanisation selects for highly mobile species with more generalist diets and reproductive strategies that are better able to exploit available resources. This syndrome was observed in bats and carabid beetles, with both groups displaying increases in traits related to mobility and generalist diets, a broader range of roosting sites for bats, and an increase in the proportion of species overwintering as adults in carabids. The shift in body size for these two taxa differed, but in ways that were consistent with increased mobility. Bats showed an increase in body size, which is consistent with previous studies that found urban environments tend to select for larger bats that are stronger and more rapid fliers, and that forage on insects in open settings using echolocation^43^. Carabids displayed a shift towards smaller bodied species^30^ that can fly^44^, a set of traits that enables greater mobility and an increased capacity to seek out food resources, without the need for strong site fidelity as observed in the central place forager or site specialist urban trait syndromes.

**Figure 5.**
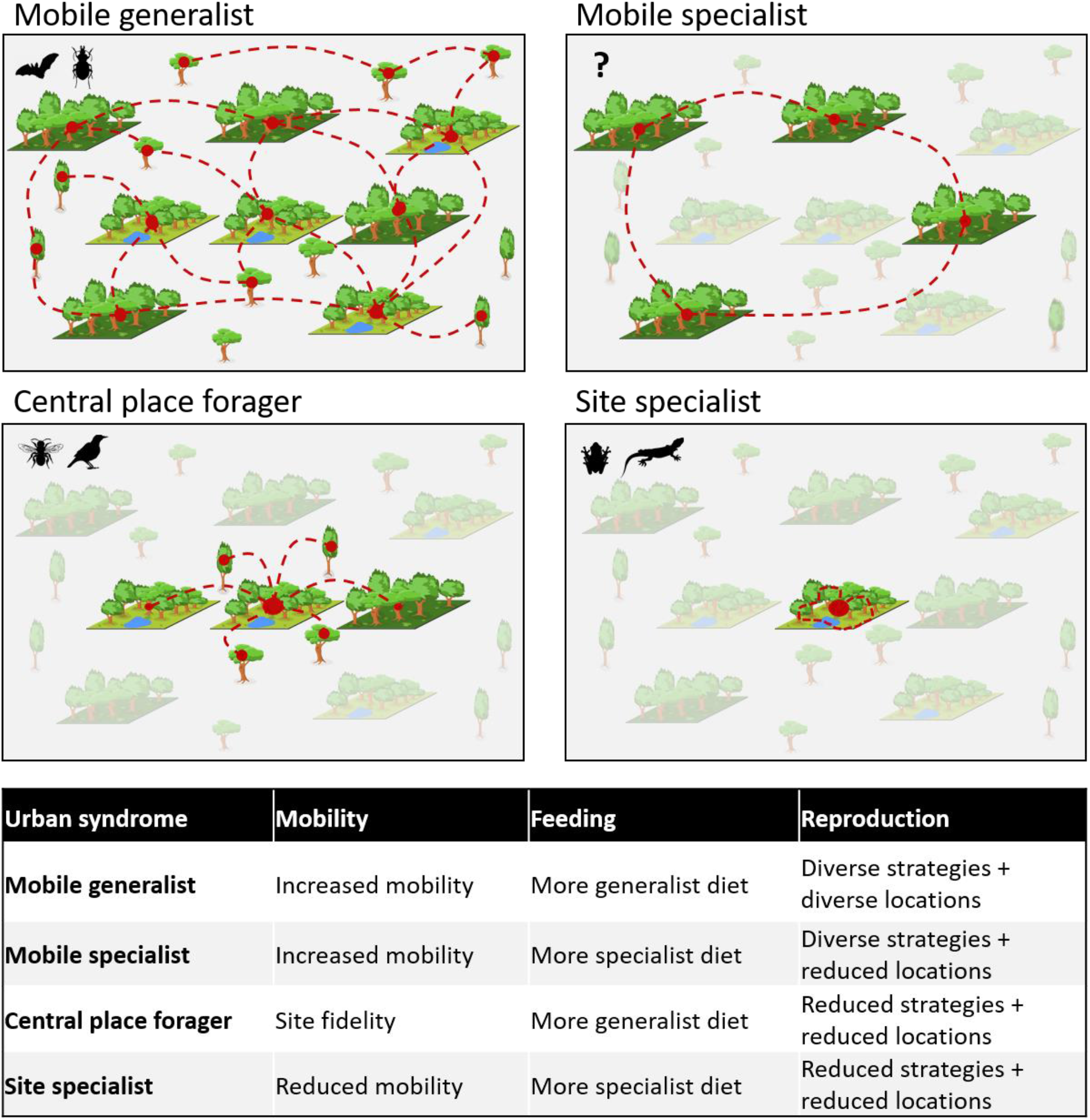
Simplified representation of the four urban trait syndromes. Two types of green habitat patches with different resources are represented in an otherwise mostly unsuitable urban matrix. Grey patches represent green habitats that are unusable for a specific taxon. Red dashed lines show typical movement pattern of taxa among patches.

The urban trait syndrome associated with site specialists was characterised by reduced mobility, increased dietary specialism and a shift towards smaller clutch sizes. All these traits are advantageous to species that are reliant on highly localised life cycles either due to resource scarcity or increased risk of mortality in the urban matrix due to predation, pollution or vehicle collision. The taxa that displayed this urban trait syndrome were amphibians and reptiles. Dietary specialisation could allow multiple species to co-exist within a more constrained physical space through resource partitioning, while reduced clutch sizes would help minimise density dependent mortality in species that are not highly mobile. Alternatively, remnant urban green spaces could act as ecological traps that disproportionally affect specialised species over generalist ones^45^, with diversity eventually decreasing as the extinction debt becomes realised^46^.

Central place foraging is an evolutionary ecology model that has been used to describe the foraging strategies for bees, mussels and other taxa^47^. As the name suggests, central place foragers establish a home base location from which they undertake daily movements to forage for additional resources. The taxa that displayed this urban trait syndrome in our study were bees and birds. Bees showed a shift towards a more solitary reproductive strategy, reduced mobility and increased dietary generalisation at very high levels of urbanisation (> 80 %, Fig. 2). For bees, this trait syndrome is consistent with previously documented movements observed in urban systems^48^. For birds, this trait syndrome was associated with reduced mobility and clutch sizes, similar to the site specialists discussed above, but accompanied here by an increase in the proportion of omnivory which would allow the individual to exploit a wider range of resources in the area surrounding their nest.

The final urban trait syndrome associated with mobile specialists is characterised by species that are able to meet their resource needs by being dietary specialists that are highly mobile and can move between spatially isolated food sources without having to return to a central place. While this urban trait syndrome was not observed in our study, there is anecdotal support for it at the species level. Wetland birds offer a useful example, where their distribution is tightly linked to a specific resource (waterbodies), but they have the capacity to easily move between locations when resources fluctuate.

While the general urban trait syndromes identified in this study are relatively clear and well supported, the associated shifts in functional diversity metrics and species richness are less consistent (Fig. 2). This may be due to differences among taxa as they relate to large-scale factors such as legacy effects that control how and to what extent regional diversity influences local diversity through species-pool effects^49^. Alternatively, if urbanisation selects for ecological strategies (or trait syndromes) that allow taxa to maximise the use of available resources, then the implications for functional diversity and species richness will be emergent properties of the species and taxonomic responses to the specifics of the resources in question. Depending on the heterogeneity and availability of resources, trait selection may result in an increase or decrease in particular trait combinations (FRic), with different levels of clustering (FEve) and expansion or contraction of the trait space (FDis). This filtering can affect community dynamics and stability through modifications of species interactions and demography^50^, and likely changes the capacity of urban biodiversity to respond to climate change and other stressors.

Our study was interested in community level trait characteristics at the taxa level. Therefore, it is quite possible that individual species within each taxon belong to different urban trait syndrome groups. For example, small insectivorous birds may display traits characteristic of site specialists, while parrots could display mobile generalist traits, and waterbirds could display mobile specialist traits. Similarly, bats are often considered to be central place foragers in other landscapes. Future research could investigate the degree to which these syndromes are representative of species within the different taxa, and how trends in functional diversity emerge from species and taxonomic responses to resource availability in urban landscapes. This information could then be used to identify resources that are critically limiting for functional diversity in urban areas and guide actions aimed at making cities suitable environments for a wider range of species.

Our results provide further evidence to counter the fallacies that urban environments are biological deserts^2^, and that biodiversity conservation is incompatible with urban areas^51^.

Instead, they point to the importance of resources, particularly those related to reproduction, as a critical filter in determining the diversity of terrestrial animals that persist in urban landscapes.

Since urbanisation occurs disproportionately in biodiversity hotspots^52^, it has been framed as a strong driver of biodiversity loss at the global scale. Our analysis shows that the diversity of species (and functional traits) found within urban areas reflects the heterogeneity and availability of resources across the urban environment. Whether populations of site scale specialists are viable or small sites are acting as ecological traps will vary on a case by case basis, particularly when supportive human actions such as ecology with cities^53^ are taken into account. Thus, our research presents a clear mandate to find innovative means of incorporating terrestrial animals’ habitat requirements (particularly related to reproductive strategies) back into cities using both land-sharing and land-sparing approaches^54^.

To maximise urban biodiversity, conservation and management should identify those species most at risk of local extinction, then determine if there are options to incorporate any limiting resources back into the landscape. However, the complexity of responses and mechanisms observed in this study suggest that positive actions for one taxon (e.g., increasing tree canopy cover for birds) may disadvantage others (such as bees that forage in more open landscapes). It follows that identifying priorities in urban biodiversity management will become an increasingly important challenge that will need to be addressed at multiple spatial scales, across diverse taxa and sites, and using a systems approach. However, the fine scale heterogeneity present in urban landscapes and the call to provide a portfolio of places to cater to diverse human preferences both offer important signals that multiple resources needs can be met within the urban landscape.

Overall, our results suggest that resource distribution and abundance are filtering taxa into one of four urban trait syndromes: **mobile generalists, mobile specialists (nomads), central place foragers and site scale specialists**. These urban trait syndromes can be applied at the level of individual species, but this study also suggests that predominant urban trait syndromes also emerge at the taxa level. Accounting for diverse urban trait syndromes and integrating them into the planning, design and management of urban environments will become increasingly critical if we are to preserve diverse biotic communities essential to the functioning of urban ecosystems and reframe the role that cities play in the global biodiversity extinction crisis.

## METHODS

### Urban biodiversity data

To identify potential datasets for our analysis, we conducted a systematic review of the published urban biodiversity literature from 1990 to 2016 to identify studies that met the following criteria: 1) community level data, 2) collected in multiple plots, and 3) across one or multiple cities. Further details about the systematic review are available in Supplementary Notes 1. Our final dataset consisted of information from 72086 plots spread across 379 cities worldwide and retained six taxonomic groups with sufficient data for a global assessment of urbanisation effects (see Fig. 1 and Supplementary Tables S1 and S2): amphibians (140 species, 1202 plots in 191 cities), bats (84 species, 540 plots in 43 cities), bees (486 species, 471 plots in 25 cities), carabid beetles (327 species, 889 plots in 17 cities), reptiles (98 species, 321 plots in 71 cities) and birds (4167 species, 68558 plots in 177 cities). The latter was collected from the eBird global community-science program (https://ebird.org)^55^, and covers the period from 1 January 2002 to 31 December 2018 from across the globe. We retained eBird checklists for analysis that were located within 1.5 km of the center of each city and were conducted using the P20, P21, P22, P23, P48, and P62 sampling protocols. We retained traveling surveys that were <1 km and area surveys that were <1 km2. We only considered observations that were identified as valid by the eBird review process, and we combined observations in grouped checklists into single checklists. While there are documented biases within this dataset^56, 57^, the signals are likely to be dampened in this study by including data points across a large number of globally distributed cities.

Within our study a plot is defined as an individual location where a survey was conducted. Therefore, while we were unable to explicitly quantify a regional species pool for each taxon and city due to limitations of the available data, we were able to quantify the level of urbanisation in the surrounding landscape for each site and confirm that our data covered the full range of values (See Fig. 2). Therefore, we are confident that our data include species outside the urban area and not simply species that are associated with urban environments.

For each taxon, we gathered functional trait data related to body size, diet, mobility and reproductive strategy, because these traits are important for an individuals’ survival, growth and reproduction^18^. We deliberately included both native and introduced species as we were interested in understanding global trait responses of species, as opposed to just the functional traits related to invasion and establishment (e.g., introduced species) or persistence and extinction risk (e.g., native species). When necessary, we standardised and simplified functional traits to ensure that the data were comparable across taxa and study areas (see Supplementary Table S3 for more detailed information; see also https://sites.rutgers.edu/urbionet/).

In addition, we analysed the community-level shifts in taxon-specific traits to account for the idiosyncrasies of each group (further details of these traits are given in Supplementary Table S3). We treated species data as presence/absence since abundance information was not available for all plots.

### Urban environment characterisation

We quantified the landscape context of each plot using data from the Global Human Settlement (GHS) images analytics framework (http://ghsl.jrc.ec.europa.eu/ghs_bu_s1.php) and the Global Forest Change database^58^. These data estimate urban extents during 2016 and forest cover during the period 2000 to 2019, thus providing a reasonable estimate of land cover, as the time ranges overlap with that of the selected studies. We included the forest cover to provide an alternative landscape to the built urban land cover, in recognition that vegetation cover can be important in driving species distributions, yet different types of vegetation offer different potential resources and habitat. We recognise that for cities in more arid landscapes, forest may not reflect the natural vegetation communities, but we consider it to still be a useful landscape type given the emphasis of urban forest strategies on increasing tree canopy cover. We calculated the percent cover and level of aggregation of urban and forest land cover within a radius of 100 m and 500 m centered on each plot for all taxa except birds, for which we use a 1000 m radius centered on each eBird checklist. We calculated the percent urban land cover in a region as the percent cover of 30 m x 30 m cells dominated by urban features (including all built-up features) using GHS. We calculated the percent forest land cover in a region with the same method, using the Global Forest Change database. To account for landscape configuration, we calculated an aggregation index^59^, which is defined as the ratio of “actual shared edges” versus “maximal possible shared edges” of the 30 m x 30 m cells. Because map units do not affect the calculation, the aggregation index can be compared among classes from the same or different landscapes and even the same landscape under different buffer sizes because the map units do not affect the calculation.

We included latitude and climate data in our analyses since the composition of functional traits have been shown to vary with latitude and climate^60, 61^. Latitude was based on the geographic coordinate of the sampling plot. The main trends in climatic conditions were characterized using the 19 Bioclim variables of the CHELSA database^62^, which provides information about biologically relevant aspects of climate for a period ranging from 1979 to 2013. We reduced the dimensionality of this dataset to limit the number of climate variables and avoid their correlations. Specifically, we ran a PCA with 100000 randomly sampled cells. We then projected the remaining cells onto the PCA. The first four PCA axes represented the main trends in climate, that is, gradients in mean temperature (PC1), diurnal range (PC2), temperature seasonality (PC3) and precipitation seasonality (PC4). Altogether, these four axes accounted for ∼89 % of the global variation in climate (see also Supplementary Table S10) and were selected for use in the subsequent analyses.

### Functional composition of animal communities

We assessed the functional composition of the species assemblage of each taxonomic group separately. This was done by calculating the community-level mean values of each trait in each plot for each taxon or, in the case of categorical traits, the proportion of species in each category. We also calculated 10 indices capturing complementary aspects of functional trait variation: functional dispersion, functional richness, and functional evenness (see Supplementary Notes 2 for further information). Since we specifically focus on functional diversity, we selected, for each aspect, the index showing the lowest correlation to species richness across all taxonomic groups (Fig. S4 Correlations among FD facets). We retained the functional dispersion (FDis), functional richness (FRic), and functional evenness (FEve) indices calculated using the alpha.fd.multidim function in the R package “mFD”^63^. Functional dispersion (FDis) measures the mean distance of individual species to the centroid of all species in multidimensional trait space^62^. A decrease in FDis shows a lower dispersion of species in trait space. FDis captures aspects of both functional richness and functional evenness. Functional richness is the amount of functional niche space occupied by species within a community^64^ and was calculated using the revised FRic index^63^. A decrease in FRic values suggests a decrease in the amount of functional trait space occupied by a community. Functional evenness measures how evenly species are distributed within the trait space (FEve index^65^). A decrease in FEve shows that species are less evenly distributed in trait space compared to the maximum possible (i.e., evenness = 1).

### Effect of urbanisation on faunal community functional composition

We analysed the global effect of urban land cover on functional community composition of each taxon while controlling for the effects of forest land cover, climatic region and latitude (see Supplementary Methods 1 for more information about the correlations among predictors).

To do so, we built various models using the random forests algorithm^66^. The random forest algorithm excels at extracting patterns from complex datasets and is becoming more common in ecological studies. This approach being nonparametric, the data need not come from a specific distribution (e.g., Gaussian) and can contain collinear variables^67^. Also, random forests can deal with model selection uncertainty because predictions are based on a consensus of many models and not just a single model selected with some measure of goodness of fit. Specifically, we used the different community functional metrics as response variables, and climate PCA axes, latitude, and the percent and aggregation of urban and forest land cover as explanatory variables. Because of the observed autocorrelation in model residuals, we added spatial covariates as explanatory variables to the models. As spatial covariates, we used positive Moran’s Eigenvector Maps of a distance matrix among sites (dbMEM)^68^. Relevant dbMEM were selected using a forward selection procedure based on the residuals of models computed without spatial covariates. The random forest algorithm was trained on 75 % of the data and evaluated on the remaining 25 %. Model training and parameter tuning were done using 2 different cross-validation strategies: 3 time 3-fold stratified CV and 30-fold spatial CV. In stratified CV, partition is stratified according to the response variable in order to balance the class distributions within the splits (function “createDataPartition” in the R package “caret”). In spatial CV, we created 30 spatial folds for cross validation (function “CreateSpacetimeFolds” in the R package “CAST”) in order to maximise the spatial transferability of model results and avoid potential overfitting. Parameter tuning used 10 random values of the number of variables to be sampled at each split time. The best model was chosen based on RMSE, MAE, and R^2^ measured on the trained dataset. The performances of the selected model were further evaluated on the test dataset using the same metrics. Spatial autocorrelation in model residuals was examined using Mantel correlograms (function “correlog” in the R package “vegan”). Potential overfitting was double-checked by comparing the model evaluation metrics among the train and test sets. We retained the models based on the spatial cross-validation procedure and including spatial covariates because they showed the overall best performances and the lowest potential overfitting and spatial autocorrelation of residuals.

To assess the importance of global drivers of changes in urban community functional composition, we estimated the importance of each explanatory variable using the residual sum of squares (RSS) from random forests models. This allowed us to assess the importance of urbanisation variables amid the influence of biogeographic and macroecological processes and determine which of latitude, climatic regions, and the percent and spatial aggregation of urban land cover induce stronger changes in community functional composition.

To assess the changes in functional community composition metrics while limiting the influence of other descriptors, we used partial dependence plots (PDP)^69, 70^. Partial dependence plots are especially useful for visualising the relationships discovered by complex machine learning algorithms such as random forests. PDPs help visualise the relationship between a subset of the features and the response while accounting for the average effect of the other predictors in the model (see Fig. 2).

All statistical analyses were performed in R version 4.0.3^71^.

## Data and code availability statement

The authors declare that the data supporting the findings of this study are available within the paper and its supplementary information files. All the code used in the analyses is open source and available in various R packages. A compiled version of the full code used for analysis is provided in a repository at https://gitlab.com/urbionet/Trait_urban_syndromes.

## Acknowledgements

This research was conducted as part of the Urban Biodiversity Research Coordination Network (UrBioNet) funded by the National Science Foundation (NSF RCN: DEB 1354676/1355151).

We initiated this project as part of the Workshop Group “Patterns, Drivers and Traits” of the Urban Biodiversity Research Coordination Network (UrBioNet, https://sites.rutgers.edu/urbionet/dsg/). We would like to thank Madhusan Katti, Christopher Trisos, and Julie Goodness for helping to conceptualise the study at the New Jersey workshop; Eliana Geretz and Carmela M. Buono for assistance with initial data compilation; and Béla Tóthmérész, Cecilia Tobar-Suárez, Etienne Normandin, Gary Luck, Lisa Smallbone, Maryna Kyrychenko-Babko, Rebecca Acosta, Yocoyani Meza-Parral for providing data included in this study.

## Authors’ contribution

The contribution of all of the people who contributed to this project is stated in the Author’s CREDIT statement, or in the Acknowledgements section of this paper. Criteria for authorship followed the Authorship Policy for UrBioNet based on Weltzin et al. (2006)^72^, and required a contribution both leading up to the analysis, and post-analysis. The process we followed for managing contributions to this project are described in Supplementary Notes 1.

## Author Contributions

Adriana Herrera-Montes: Conceptualization (NJ workshop); Methodology (NJ Workshop); Data Curation (Taxa coordinator- Reptiles/Amphibians); Writing – review & editing (round 1 results); Writing – review & editing (final polished draft ms); Writing – review & editing (final Version 2 ms)

Alan Vergnes: Data Contributor; Writing – review & editing (round 1 results); Writing – review & editing (final Version 2 ms)

Aldemar A. Acevedo: Data Contributor; Writing – review & editing (round 1 results); Writing – review & editing (final Version 2 ms)

Alison M. Gainsbury: Data Contributor; Writing – review & editing (round 1 results); Writing – review & editing (final Version 2 ms)

Allyson B. Salisbury: Conceptualization (NJ workshop); Methodology (NJ Workshop); Data curation; Formal Analysis; Visualisation; Writing – review & editing (round 1 results); Writing – review & editing (final polished draft ms); Writing – review & editing (final Version 2 ms)

Amy K. Hahs: Conceptualization (RCN proposal and NJ workshop); Methodology (NJ Workshop); Project Administration; Supervision; Data Curation (Spatial data); Visualisation; Writing – original draft; Writing – review & editing (round 1 results); Writing – review & editing (final polished draft ms); Writing – drafting, review and editing Version 2 and response to reviewers comments

Ana Rainho: Data Contributor; Writing – review & editing (full manuscript draft); Writing – review & editing (final Version 2 ms)

Andrew J. Hamer: Data Contributor; Writing – review & editing (round 1 results); Writing – review & editing (final Version 2 ms)

Assaf Schwartz: Data Contributor; Writing – review & editing (full manuscript draft); Writing – review & editing (final Version 2 ms)

Bertrand Fournier: Methodology; Data Analyses; Curation (polishing and matching datasets); Visualisation (original and Version 2); Writing – original draft; Writing – review & editing (round 1 results); Writing – review & editing (final polished draft ms); Data Analysis Version 2; Writing – drafting, review and editing Version 2 and response to reviewers comments

Caragh Threlfall: Conceptualization (NJ workshop); Methodology (NJ Workshop); Data Curation (Taxa coordinator- Bees, Bats); Writing – review & editing (round 1 results); Writing – review & editing (final polished draft ms); Writing – review & editing (final Version 2 ms)

Charles H. Nilon: Conceptualization (RCN proposal and NJ workshop); Methodology (NJ Workshop); Funding; Writing – review & editing (round 1 results); Writing – review & editing (final Version 2 ms)

Christian Voigt: Data Contributor; Writing – review & editing (full manuscript draft)

Christine Rega-Brodsky: Data Curation (Taxa coordinator – Birds); Writing – review & editing (round 1 results); Writing – review & editing (final polished draft ms); Writing – review & editing (final Version 2 ms)

Christopher A. Lepczyk: Conceptualization (RCN proposal and NJ workshop); Methodology (NJ Workshop); Writing – review & editing (round 1 results); Writing – review & editing (final polished draft ms); Writing – review & editing (final Version 2 ms)

D. Johan Kotze: Conceptualization (NJ workshop); Methodology (NJ Workshop); Supervision; Data Curation (Taxa coordinator- Carabids); Writing – original draft; Writing – review & editing (round 1 results); Writing – review & editing (final polished draft ms); Writing – drafting Version 2 and response to reviewers comments

Daniel Lewanzik: Data Contributor; Writing – review & editing (full manuscript draft); Writing – review & editing (final Version 2 ms)

David M. Lowenstein: Data Contributor; Writing – review & editing (round 1 results); Writing – review & editing (final Version 2 ms)

David O’Brien: Data Contributor; Writing – review & editing (round 1 results); Writing – review & editing (final Version 2 ms)

Desiree Tomassi: Data Contributor; Writing – review & editing (round 1 results)

Eduardo Pineda: Data Contributor; Writing – review & editing (full manuscript draft); Writing – review & editing (final Version 2 ms)

Ela S. Carpenter: Conceptualization (NJ workshop); Methodology (NJ Workshop); Data Contributor; Writing – review & editing (full manuscript draft)

Elena Belskaya: Data Contributor; Writing – review & editing (round 1 results); Writing – review & editing (final Version 2 ms)

Frank A. La Sorte: Conceptualization (RCN proposal and NJ workshop); Methodology (NJ Workshop); Data curation; Software; Formal Analysis; Visualisation; Writing – review & editing (round 1 results); Writing – review & editing (final polished draft ms); Writing – review & editing (final Version 2 ms)

Gabor Lövei: Data Contributor; Writing – review & editing (round 1 results)

Ian MacGregor-Fors: Conceptualization (NJ workshop); Methodology (NJ Workshop); Data Curation (Taxa coordinator- Reptiles/Amphibians); Writing – review & editing (round 1 results); Writing – review & editing (final polished draft ms); Writing – review & editing (final Version 2 ms)

J. Scott MacIvor: Conceptualization (NJ workshop); Methodology (NJ Workshop); Data Curation (Taxa coordinator- Bees); Writing – review & editing (round 1 results); Writing – review & editing (final polished draft ms); Writing – review & editing (final Version 2 ms)

James C. Makinson: Data Contributor; Writing – review & editing (round 1 results); Writing – review & editing (final Version 2 ms)

Jennifer Castañeda-Oviedo: Data Contributor; Writing – review & editing (round 1 results)

Joanna Coleman: Data Contributor; Writing – review & editing (full manuscript draft); Writing – review & editing (final Version 2 ms)

Jon P. Sadler: Data Contributor; Writing – review & editing (round 1 results); Writing – review & editing (final Version 2 ms)

Jordan Shroyer: Data Contributor; Writing – review & editing (round 1 results)

Julie Teresa Shapiro: Data Contributor; Writing – review & editing (full manuscript draft); Writing – review & editing (final Version 2 ms)

Katherine C.R. Baldock: Data Contributor; Writing – review & editing (round 1 results); Writing – review & editing (full manuscript draft); Writing – review & editing (final Version 2 ms)

Kelly Ksiazek-Mikenas: Data Contributor; Writing – review & editing (round 1 results); Writing – review & editing (final Version 2 ms)

Kevin C. Matteson: Data Contributor; Writing – review & editing (round 1 results); Writing – review & editing (final Version 2 ms)

Kirsten Jung: Conceptualization (NJ workshop); Methodology (NJ Workshop); Data Curation (Taxa coordinator- Bats); Writing – review & editing (round 1 results); Writing – review & editing (final polished draft ms); Writing – review & editing (final Version 2 ms)

Kyle Barrett: Data Contributor; Writing – review & editing (round 1 results)

Lizette Siles: Data Contributor; Writing – review & editing (round 1 results); Writing – review & editing (final Version 2 ms)

Luis F. Aguirre: Data Contributor; Writing – review & editing (round 1 results); Writing – review & editing (final Version 2 ms)

Luis Orlando Armesto: Data Contributor; Writing – review & editing (round 1 results)

Marcin Zalewski: Data Contributor; Writing – review & editing (round 1 results); Writing – review & editing (final Version 2 ms)

Marco Moretti: Conceptualization (NJ workshop); Methodology (NJ Workshop); Supervision; Data Curation (Taxa coordinator- Carabids); Writing – original draft; Writing – review & editing (round 1 results); Writing – review & editing (final polished draft ms); Writing – drafting Version 2 and response to reviewers comments

Maria Isabel Herrera-Montes: Data Contributor; Writing – review & editing (round 1 results); Writing – review & editing (final Version 2 ms)

Martin K. Obrist: Data Contributor; Writing – review & editing (round 1 results)

Max R. Piana: Conceptualization (NJ workshop); Methodology (NJ Workshop); Data curation; Writing – review & editing (round 1 results); Writing -review & editing (final polished draft ms)

Myla F.J. Aronson: Conceptualization (RCN proposal and NJ workshop); Methodology (NJ Workshop); Funding; Writing – review & editing (round 1 results); Writing – review & editing (final polished draft ms); Writing – review & editing (final Version 2 ms)

Nicholas S.G. Williams: Conceptualization (RCN proposal and NJ workshop); Methodology (NJ Workshop); Data Curation (Taxa coordinator- Bees); Writing – review & editing (round 1 results); Writing – review & editing (final polished draft ms); Writing – drafting (final Version 2 ms)

Rebecca K. Tonietto: Data Contributor; Writing – review & editing (round 1 results) Ricardo Torrado: Data Contributor; Writing – review & editing (round 1 results)

Sara A. Gagne: Data Contributor; Writing – review & editing (round 1 results); Writing – review & editing (final Version 2 ms)

Sarah J. Hinners: Data Contributor; Writing – review & editing (round 1 results); Writing – review & editing (final Version 2 ms)

Sonja Knapp: Conceptualization (NJ workshop); Methodology (NJ Workshop);Data Curation (Taxa coordinator- Reptiles/Amphibians); Writing – review & editing (round 1 results); Writing – review & editing (final polished draft ms); Writing – review & editing (final Version 2 ms)

Tanya Latty: Data Contributor; Writing – review & editing (round 1 results)

Thilina D. Surasinghe: Data Contributor; Writing – review & editing (round 1 results); Writing – review & editing (final Version 2 ms)

Thomas Sattler: Data Contributor; Data curation (Birds); Writing – review & editing (round 1 results); Writing – review & editing (final Version 2 ms)

Tibor Magura: Data Contributor; Writing – review & editing (round 1 results); Writing – review & editing (final Version 2 ms)

Werner Ulrich: Data Contributor; Writing – review & editing (full manuscript draft); Writing – review & editing (final Version 2 ms)

Zoltan Elek: Data Contributor; Writing – review & editing (full manuscript draft); Writing – review & editing (final Version 2 ms)

