## Supplementary information for "Urbanisation generates multiple trait syndromes for terrestrial taxa worldwide"

**Institutes:** [1]School of Ecosystem and Forest Sciences, The University of Melbourne, Burnley Campus 500 Yarra Blvd, Richmond 3121 VIC Australia ; [2]Institute of Environmental Science and Geography, University of Potsdam, Karl-Liebknecht-Str. 24-25, 14476 Potsdam, Germany; [3]Department of Ecology, Evolution and Natural Resources, Rutgers, The State University of New Jersey, New Brunswick, NJ 08816 USA, ; [4]School of Natural Resources, University of Missouri, Columbia, MO 65211 USA; [5]Department of Environmental Science, College of Natural Sciences, University of Puerto Rico, ; [6]The Morton Arboretum, 4100

Illinois Route 53, Lisle, IL 60532, USA ; [7]School of Life and Environmental Sciences, The University of Sydney, NSW 2006, Australia; [8]Department of Biology, Pittsburg State University, Pittsburg, KS 66762 USA; [9]School of Forestry and Wildlife Sciences, Auburn University, Auburn, AL 36849, USA; [10]U.S. Fish and Wildlife Service, Chesapeake Bay Field Office, 177 Admiral Cochrane Dr. Annapolis, MD 21401; [11]Cornell Lab of Ornithology, Cornell University, Ithaca, NY, 14850 USA; [12]Faculty of Biological and Environmental Sciences, Ecosystems and Environment Research Programme, Niemenkatu 73, FI-15140, Lahti, University of Helsinki, Finland; [13]Department of Biological Sciences, University of Toronto Scarborough, 1265 Military Trail, Toronto Canada M1C 1A4; [14]Institute of Evolutionary Ecology and Conservation Genomics, Ulm University, Albert-Einstein-Allee 11, 89069 Ulm, Germany; [15]USDA Forest Service, Northern Research Station, Amherst, MA 01002 USA ; [16]Helmholtz Centre for Environmental Research – UFZ, Department of Community Ecology, Theodor-Lieser-Str. 4, 06120 Halle (Saale), Germany ; [17]German Centre for Integrative Biodiversity Research (iDiv) Halle-Jena-Leipzig, Puschstraße 4, 04103 Leipzig, Germany ; [18]Department of Ecology, Ecosystem Science/Plant Ecology, Technische Universität Berlin, 12165 Berlin, Germany ; [19]CEFE, Univ Montpellier, CNRS, EPHE, IRD, Univ Paul Valéry Montpellier 3, Montpellier, France; [20]Departamento de Ciencias Ecológicas, Facultad de Ciencias, Laboratorio de Genética y Evolución, Universidad de Chile. Las Palmeras 3425, Ñuñoa, Santiago, Chile; [21]University of South Florida, St. Petersburg Campus, Department of Integrative Biology, St. Petersburg, FL, 33701, USA; [22]Centre for Ecology, Evolution and Environmental Changes (cE3c) and Dept. of Animal Biology, Faculty of Sciences, Univ. of Lisbon, Lisboa, Portugal; [23]Institute of Aquatic Ecology, Centre for Ecological Research, Karolina u. 29, 1113 Budapest, Hungary; [24]Faculty of Architecture and Town Planning, Technion – Israel Institute of Technology, Haifa, 32000, Israel; [25]Dept. of Evolutionary Ecology, Leibniz Institute for Zoo and Wildlife Research, Alfred-Kowalke-Str.

17, 10315 Berlin, Germany; [26]Michigan State University Extension, Macomb County, MI, USA; [27]Scottish Natural Heritage (NatureScot), Great Glen House, Inverness, IV3 8NW, UK; [28]Institute of Marine Sciences, University of California Santa Cruz, Santa Cruz, CA 95064, USA; [29]Red de Biología y Conservación de Vertebrados. Instituto de Ecología, A.C. Carretera Antigua a Coatepec 351, Xalapa, 91073, Mexico; [30]Institute of Plant and Animal Ecology, Ural Branch, Russian Academy of Sciences, Eighth March Street 202, Yekaterinburg 620144, Russia; [31]Department of Agroecology, Aarhus University, DK-4200 Slagelse, Denmark; [32]Hawkesbury Institute for the Environment, Western Sydney University, Locked Bag 1797, Penrith, NSW, 2751, Australia; [33]Grupo de Investigación en Ecología y Biogeografía, Universidad de Pamplona, Pamplona, Colombia; [34]Queens College at the City University of New York, Flushing NY USA; [35]School of Geography, Earth and Environmental Sciences, University of Birmingham, Edgbaston, Birmingham B15 2TT, UK; [36]University of Lyon, French Agency for Food, Environmental and Occupational Health & Safety (ANSES), Laboratory of Lyon, 31 Avenue Tony Garnier, 69364, Lyon Cedex 07, France; [37]Department of Geography and Environment Sciences, Northumbria University, Newcastle upon Tyne, UK; [38]School of Biological Sciences, University of Bristol, Bristol, UK; [39]Cabot Institute, University of Bristol, Bristol, UK; [40]Department of Biology, Elmhurst University, Elmhurst, IL 60126 USA; [41]Department of Biology/Project Dragonfly, Miami University, Oxford, OH, USA; [42]Department of Forestry and Environmental Conservation, 261 Lehotsky Hall, Clemson University, Clemson, SC 29631; [43]Área de Mastozoología, Museo de Historia Natural Alcide d'Orbigny. Avenida Potosí 1458, Cochabamba. Cochabamba, Bolivia; [44]Centro de Biodiversidad y Genética, Universidad Mayor de San Simón, c Sucre, frente Parque La Torre s/n, Bolivia; [46]Museum and Institute of Zoology of the Polish Academy of Sciences, Wilcza 64, Warsaw 00-679, Poland; [47]Grupo de Ecologia Animal, Universidad del Valle, Cali, Colombia; [48]Swiss Federal Institute for Forest, Snow and Landscape Research

WSL, Biodiversity and Conservation Biology, CH-8903 Birmensdorf, Switzerland; [49]Department of Natural Sciences, University of Michigan-Flint, 303 E Kearsley St., Flint, Michigan, 48502, USA; [50]Secretaría de Educación del Municipio de Cúcuta, Colombia; [51]University of North Carolina at Charlotte, 9201 University City Blvd., Charlotte, North Carolina, USA, 28223; [52]Department of City and Metropolitan Planning, University of Utah, Salt Lake City, Utah, USA; [53]Sydney Institute of Agriculture. School of Life and Environmental Sciences; University of Sydney, Sydney Australia; [54]Department of Biological Sciences, Bridgewater State University, Bridgewater, MA 02325; [55]Swiss Ornithological Institute, Seerose 1, CH-6204 Sempach, Switzerland; [56]Department of Ecology, Faculty of Science and Technology, University of Debrecen, H-4032 Debrecen, Egyetem square 1., Hungary; [57]ELKH-DE Anthropocene Ecology Research Group, University of Debrecen, H-4032 Debrecen, Egyetem square 1. , Hungary; [58]Department of Ecology and Biogeography, Nicolaus Copernicus University, Lwowska 1, 87-100 Torun, Poland; [59]Centre for Agricultural Research, Plant Protection Institute, Eötvös Loránd Research Network, Herman Ottó út 15, Budapest 1022, Hungary; [60]Swiss Federal Research Institute WSL, Biodiversity and Conservation Biology, Zürcherstrasse 111, 8903 Birmensdorf, Switzerland

**Supplementary Information for** *Urbanisation generates multiple trait syndromes for terrestrial taxa worldwide*

### Table of Contents

*Supplementary Notes 1:* Additional information about data collection and trait selection

*Supplementary Notes 2:* Additional information about the computation and selection of functional diversity metrics

*Supplementary Figure S1:* Global distribution of data included in this study

*Supplementary Figure S2:* Additional information about functional diversity indices

*Supplementary Figure S3:* Species accumulation curves for each taxonomic group

*Supplementary Figure S4:* Correlations of diversity metrics across taxonomic groups

*Supplementary Figure S5:* Predicted changes in species richness values per taxon along an urbanisation gradient

*Supplementary Figure S6:* Spatial autocorrelation of model residuals using Moran's I at various distance thresholds

*Supplementary Table S1:* Summary of the dataset used in the analyses

*Supplementary Table S2:* Summary of number of cities with different numbers of taxa sampled

*Supplementary Table S3:* Additional information about traits

*Supplementary Table S4:* Detailed description of **Amphibian** traits.

*Supplementary Table S5:* Detailed description of **Bat** traits.

*Supplementary Table S6:* Detailed description of **Bee** traits.

*Supplementary Table S7:* Detailed description of **Bird** traits.

*Supplementary Table S8:* Detailed description of **Carabid beetle** traits.

*Supplementary Table S9:* Detailed description of **Reptile** traits.

*Supplementary Table S10:* Factor loadings on global climate PCA axes

*Supplementary Methods 1:* Correlations among environmental variables

*Supplementary Data 1*. Excel file with compiled dataset used for these analysis – Amphibians

*Supplementary Data 2.* Excel file with compiled dataset used for these analysis – Bats

*Supplementary Data 3*. Excel file with compiled dataset used for these analysis – Bees

*Supplementary Data 4.* Excel file with compiled dataset used for these analysis – Carabids

*Supplementary Data 5.* Excel file with compiled dataset used for these analysis – Reptiles

*Supplementary Data 6.* Excel file containing the localities used to extract bird data from eBird, and the list of bird species included in our analysis. The full Excel file with compiled Bird dataset used for these analysis has not been provided due to file size limits (it is 700MB in size), but can be extracted from eBird or obtained directly from the corresponding author.

***Supplementary Notes 1:* Additional information about data collection and trait selection**

The Web of Science literature review was conducted using the following keywords:

**TOPIC:** (urban OR “peri-urban” OR periurban* OR suburban* OR “sub-urban*” OR conurbation OR city OR cities OR town* OR megalopol* OR metropol* OR “built-up” OR “built environm*”)

*AND*

**TOPIC:** (Arthropod* OR Invertebrate* OR Insect*)

**TOPIC:** (Bats OR Chiroptera)

**TOPIC:** (Bees OR Hymenoptera OR Aculeata OR Pollinator*)

**TOPIC:** (Carabid* OR “ground beetle*” OR Cincidel* OR “tiger beetle*”)

**TOPIC:** (Mammal* OR Marsupial*)

**TOPIC:** (Amphib* OR frog* OR toad* OR Anura OR Reptil*)

*AND*

**Refined by: TOPIC:** (Biodiversity OR “species composition” OR “species assembl*” OR “species communit*” OR “species richness” OR “species diversity” OR “Shannon-Weaver” OR “Shannon-Weiner” OR “Shannon-Wiener” OR “Shannon* H” OR “Simpson* index” OR “Simpson* diversity” OR “Simpson* dissimilarity” OR “Simpson* beta” OR evenness OR “alpha diversity” OR “beta diversity” Biodiversity OR composition OR assembl* OR communit* OR richness OR diversity OR Shannon OR Simpson OR evenness) AND **TOPIC:** (Gradient* OR Grid OR Site OR Map* OR Spatial* OR GIS OR Plot OR Sample OR Area OR Occurrence OR Presence OR distribution) AND **WEB OF SCIENCE CATEGORIES:** (

ECOLOGY OR BIODIVERSITY CONSERVATION OR PLANT SCIENCES OR URBAN STUDIES OR FORESTRY )

Indexes=SCI-EXPANDED Timespan=1990-2016

It identified the published studies that had datasets suitable for our intended analysis as they contained 1) community level data, 2) collected in multiple plots, and 3) across one or multiple cities. From these studies, we identified those taxa with sufficient number of papers to permit a meta-analysis (e.g., in 2016 there were only 8 papers investigating snails in urban landscapes that met our criteria), and the methods we proposed to use would be appropriate (e.g., fish are unlikely to be strongly influenced by % tree cover within 100 – 500 m).

Once we had finalised our taxa groups, we identified a core group of authors from the WoS literature review who had datasets that would be suitable for our proposed analysis, and contacted them to see if they would be interested in collaborating in this research. We also gained additional collaborators who volunteered their datasets after hearing our presentations on this project at the 2017 Ecological Society of America meeting, 2017 Ecological Society of Australia meeting and 2017 International Ecology Congress (INTECOL).

Researchers who responded to our invitation were provided with a data collection template to ensure we received all of the required information in a format that could readily be integrated into a larger dataset. This included a species x site table, species x trait table and site information table, and a link to a metadata form where we could capture additional information about the study used to produce the data.

To facilitate this process, we formed taxa-based groups which were coordinated by 2-3 members of the UrBioNet coordinating group. Throughout the project the taxa coordinators were responsible for collating the data into a single dataset, and worked with the data contributors to develop the compiled species x trait tables. Once the data had been compiled the coordinating group populated the standardised site information, analysed the data and created a preliminary results document that was circulated back to the data contributors for discussion and feedback within the taxonomic groups in March 2020. Feedback from this process was compiled back by the coordinating group and used to update the analysis. The coordinating group then drafted a manuscript that was shared back to the data contributors for feedback. After the feedback on the manuscript had been incorporated, a final version of the manuscript was circulated to ensure all named authors agreed to the submission of the manuscript. As the timing of the circulation of the initial round of results coincided with the onset of the COVID-19 global pandemic, the timeframes for delivering this project were disrupted, and some of the additional feedback rounds were bypassed as a large number of contributors had already met the criteria for authorship.

The exception to this process was the bird taxon, where we extracted data from existing global datasets to match the cities where we had information for other taxa groups.

As this research was hosted by the UrBioNet Research Coordination Network, we applied their Authorship Policy (https://sites.rutgers.edu/urbionet/about/authorship-guidelines/), where authorship required a substantial contribution beyond simply providing data or being present at the initial workshop. This is in alignment with Weltzin et al. (2006)[71] and other publications that seek to ensure authorship reflects a substantial contribution.

***Supplementary Notes 2:* Additional information about the computation and selection of functional diversity metrics**

We first imputed missing trait values using the K-nearest neighbors (function "preProcess" in the R package "caret"). We then calculated the Gower functional distance among species based on centered and scaled trait values. We computed a Principal Coordinates Analysis (PcoA) using the resulting functional distance matrices. The quality of trait spaces was evaluated as the absolute deviation between trait-based distance and distance in the PcoA-based space[70]. The number of axes producing the lower deviation was retained for further analyses. In the case of amphibian and reptile, many sites had 3 species or less. As a result, we retained 2 PcoA axes to be able to compute functional diversity metrics for a maximum number of sites. This represented a fair compromise between the quality of the trait space (third-best option for amphibians and second-best option for reptiles) and the number of sites to be included in the analyses. Overall, we included between 48 % and 61 % of the total variation in trait data in our analyses (Amphibian = 48 % over 2 PcoA axes, Bat = 59 % over three PcoA axes, Bee = 60 % over three PcoA axes, Bird = 61 % over three PcoA axes, Carabid beetles = 61 % over three PcoA axes, Reptile = 53 % over two PcoA axes). We finally computed FDis, FRic, and FEve using the function alpha.fd.multidim of the R package "mFD"[61].

In addition, we computed the functional alpha diversity applied to distance between species[71] (function "alpha.fd.hill" of the R package "mFD"), the Rao functional dispersion index[72]; the functional dispersion index based on the "classic" framework of Laliberté & Legendre[62] (function "dbFD" of the R package "FD"), the "TOP" functional richness[73]; the "TED" functional evenness[73]; and "Fever" functional evenness[74]. An example script for the calculations of these metrics is provided in a repository at https://gitlab.com/urbionet/Trait_urban_syndromes.

61 Magneville, C. *et al.* mFD: an R package to compute and illustrate the multiple facets of functional diversity. *Ecography* **2022**, doi:https://doi.org/10.1111/ecog.05904 (2022).

62 Laliberté, E. & Legendre, P. A distance-based framework for measuring functional diversity from multiple traits. *Ecology* **91**, 299-305, doi:doi:10.1890/08-2244.1 (2010).

70 Maire, E., Grenouillet, G., Brosse, S. & Villéger, S. How many dimensions are needed to accurately assess functional diversity? A pragmatic approach for assessing the quality of functional spaces. *Global Ecol. Biogeogr.* **24**, 728-740, doi:https://doi.org/10.1111/geb.12299 (2015).

71 Chao, A. *et al.* An attribute-diversity approach to functional diversity, functional beta diversity, and related (dis) similarity measures. *Ecol. Monogr.* **89**, e01343 (2019).

72 Ricotta, C. & Moretti, M. CWM and Rao’s quadratic diversity: a unified framework for functional ecology. *Oecologia* **167**, 181-188 (2011).

73 Fontana, S., Petchey, O. L. & Pomati, F. Individual-level trait diversity concepts and indices to comprehensively describe community change in multidimensional trait space. *Funct. Ecol.* **30**, 808-818, doi:10.1111/1365-2435.12551 (2016).

74 Ricotta, C., Bacaro, G. & Moretti, M. A New Measure of Functional Evenness and Some of Its Properties. *PLoS ONE* **9**, e104060, doi:10.1371/journal.pone.0104060 (2014).

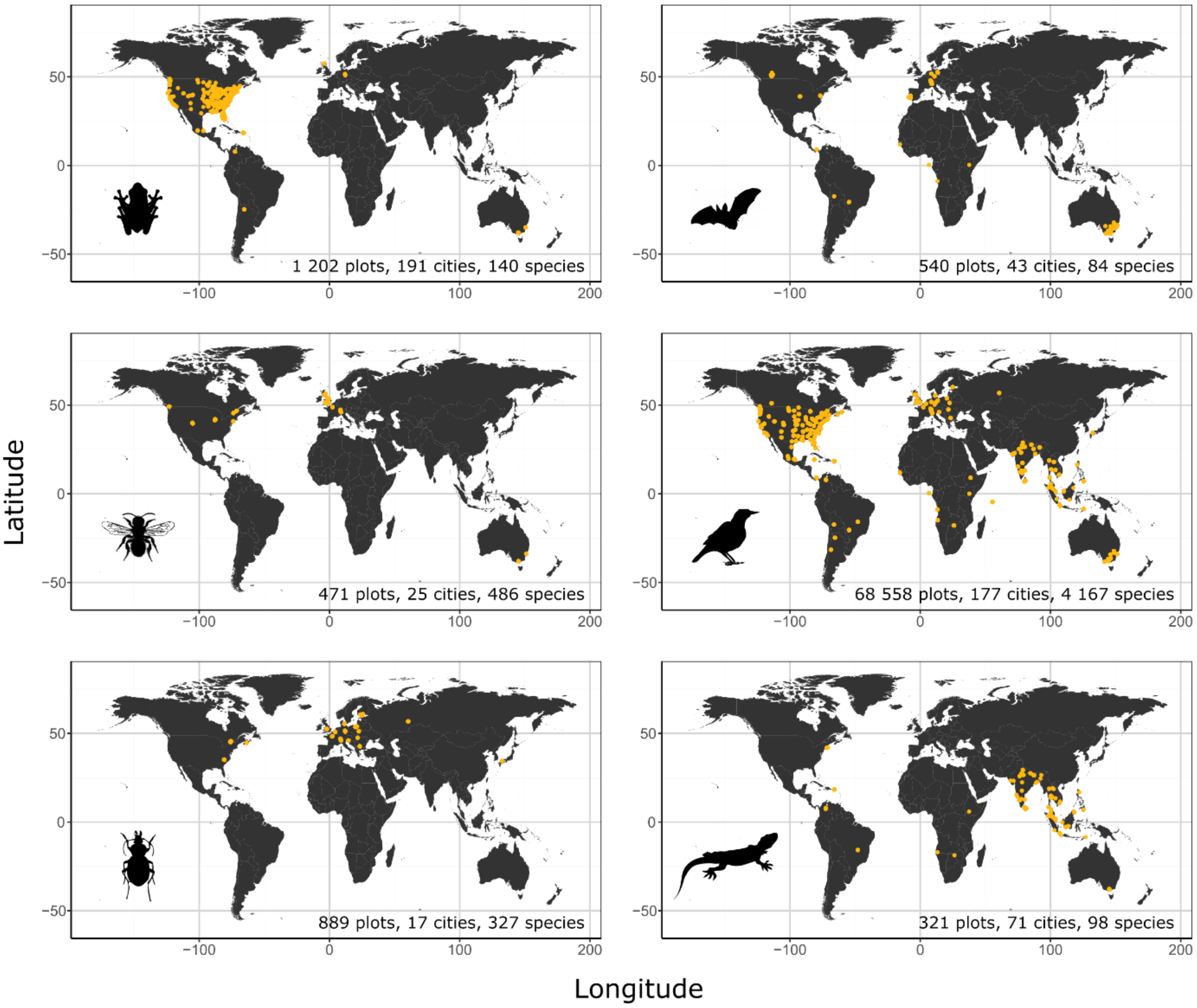

***Supplementary Figure S1:*** Global distribution of data included in this study. Locations of cities with sampling plots for each taxonomic group individually. All data come from the UrBioNet contributor network except for birds (eBird).

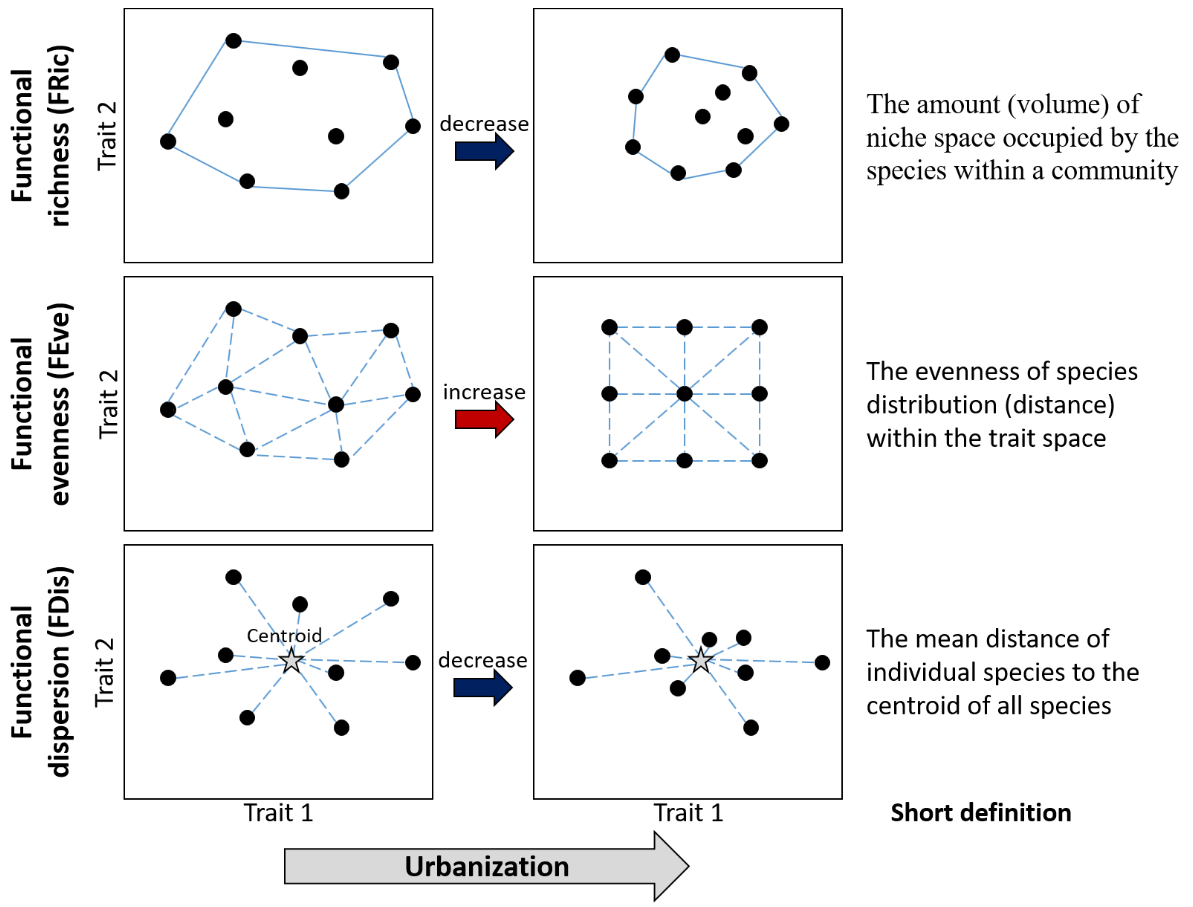

***Supplementary Figure S2.*** Additional information about functional diversity indices. Expected responses of functional richness (FRic), functional evenness (FEve) and functional dispersion (FDis) to increased urbanisation. Functional richness is expected to decrease as a result of the loss of some functional groups (environmental filtering). Functional evenness is expected to increase as a result of increased competition for more scarce resources (competitive exclusion of functionally similar species). Functional dispersion is expected to decrease because increased urbanisation is expected to select for generalist species with broad environmental tolerances (species close to the centroid). The right column provides a short definition of each index.

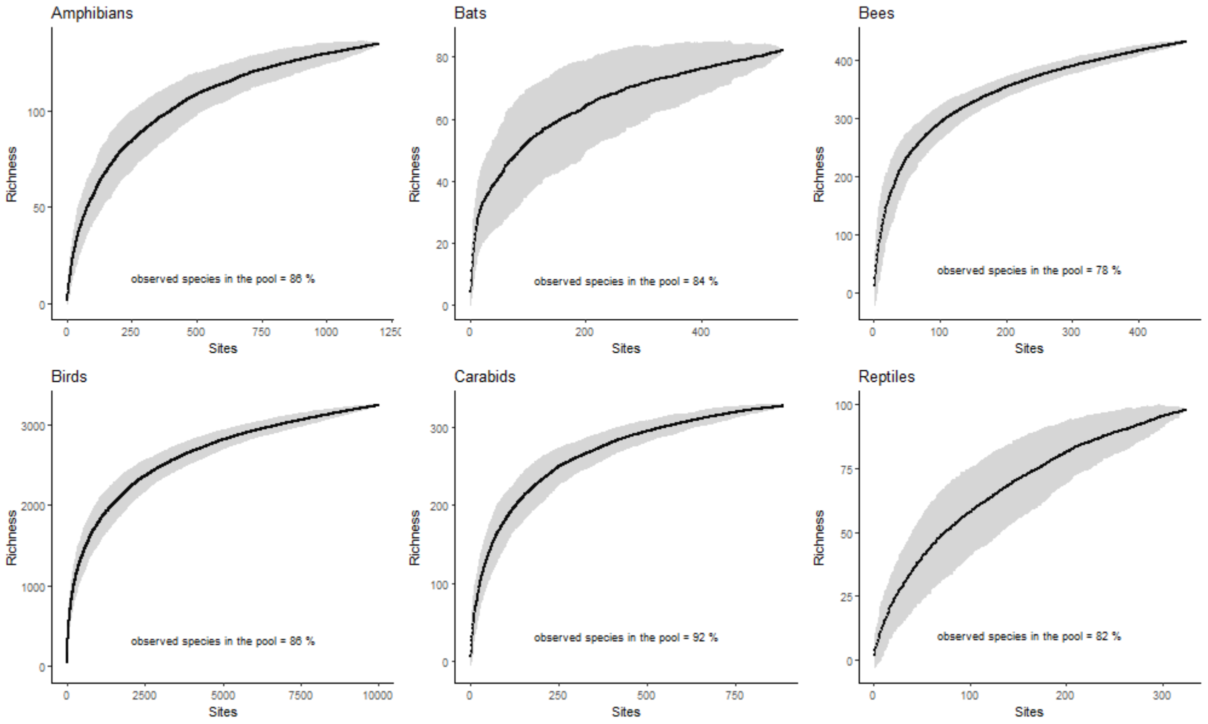

***Supplementary Figure S3***. Species accumulation curves for each taxonomic group. These curves were used to estimate the total number of species present in the global species pool (extrapolated species richness in the species pool based on bootstrap resampling).

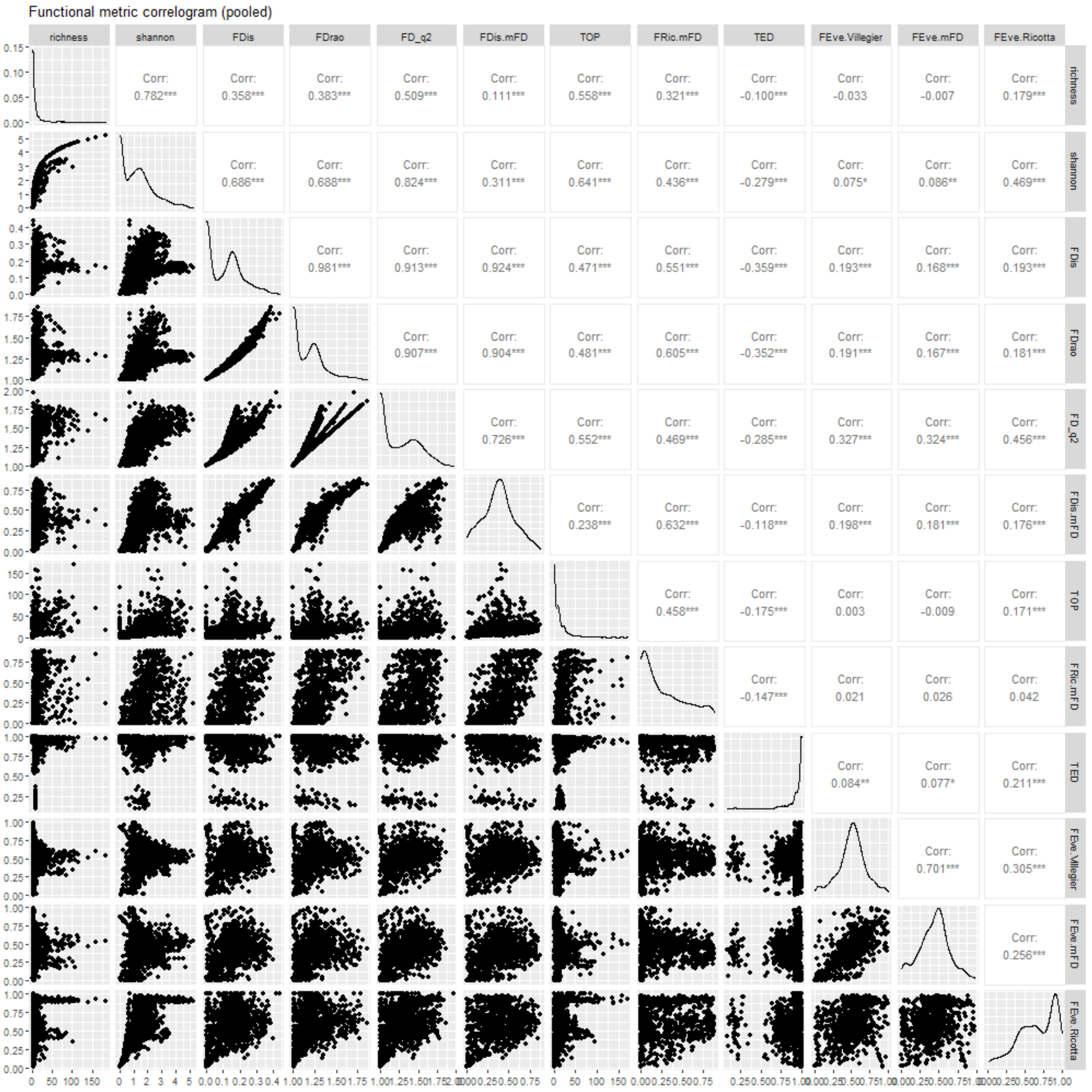

***Supplementary Figure S4***. Correlations of diversity metrics across taxonomic groups. For each taxonomic group, 300 sites were randomly chosen and pooled to ensure that each group contributes equally to this analysis. For each functional diversity facet of interest, we selected the metric showing the lowest correlations to species richness (functional dispersion = FDis_mFD, richness = FRic_mFD, evenness = FEve_mFD).

***Supplementary Table S1:*** Summary of the dataset used in the analyses and whether the data were compiled from directly contributed datasets, or e-Bird. The geographical distribution of the sampling plots for each taxonomic group is shown in Fig. 1 and Supplementary Figure S1.

| ***Taxa*** | ***N. Plots*** | ***N. Cities*** | ***N. Species*** | ***Source*** |
|---|---|---|---|---|
| Amphibians | 1 202 | 191 | 140 | UrBioNet contributor network |
| Bats | 540 | 43 | 84 | UrBioNet contributor network |
| Bees | 471 | 25 | 486 | UrBioNet contributor network |
| Birds | 68 558 | 177 | 4 167 | e-Bird |
| Carabids | 882 | 17 | 327 | UrBioNet contributor network |
| Reptiles | 324 | 71 | 98 | UrBioNet contributor network |

***Supplementary Table S2:*** Summary of number of cities with different numbers of taxa sampled. For example, only one city has been sampled for 5 taxa (Melbourne, Australia), and 3 cities have been sampled for 4 taxa (Lugano, Luzern and Zürich, Switzerland). The geographical distribution of the sampling plots for each taxonomic group is shown in Fig. 1 and Supplementary Figure S1.

| Number Taxa Sampled in the City | Number of Cities |
|---|---|
| 1 | 254 |
| 2 | 109 |
| 3 | 12 |
| 4 | 3 |
| 5 | 1 |

***Supplementary Table S3:*** Information about the evaluated traits presented in Fig. 2 of the manuscript. Specific traits are presented here but have not been shown in Fig. 2 of the manuscript. Further information around these traits and the data sources used for each individual taxonomic group can be found in Supplementary Tables S4-S9.

| ***Taxonomic group*** | ***Body size*** | ***Feeding*** | ***Mobility*** | ***Reproductive Strategy*** | ***Specific traits*** |
|---|---|---|---|---|---|
| Amphibians | **Body length** [cm] | **Diet breadth** (specialist=0; generalist=1) | **Movement Distances** (reduced=0; moderate=1; high=2) | **Clutch size** (0=small; 1 = intermediate; 2=large) | Aquatic habitat affinity index (0=low;1=medium; 2=high) |
| Bats | **Forearm length** [mm] | **Hunting strategy** (gleaning=1; others=0) | **Aspect ratio** | **Roosting requirements** (specialist=0; generalist=1) | Wing loading (nb) / Echolocation (kHz) / Dispersal strategy (mobility in open habitats=1; others=0) |
| Bees | **Inter-tegula distance** [mm] | **Tongue length** (short tongue=1; long tongue=0) | **Inter-tegula distance** [mm] | **Sociality** (Solitary =1; other=0) | Nesting strategy (Below ground (Below ground =1; others=0) / Above ground (Above ground =1; others=0) / Parasite (Parasite=1; others=0)) |

| | | | | | |
|---|---|---|---|---|---|
| Birds | **Body mass** [g] | Trophic niche (**omnivorous**=1; others=0 / Fruit-nectar=1; others=0 / Invertebrate=1; others=0 / Plant-seed=1; others=0 / Vertebrates-scavenger =1; others=0) | **Hand-wing index** | **Clutch size** (number of eggs) | Foraging strata index (Habitat [0=aquatic; 1=terrestrial; 2=aerial]; Aquatic [0-2]; Terrestrial [0-4], Aerial [0-1]) |
| Carabids | **Body length** [cm] | Trophic guild (Herbivore=1; others=0, Carnivore= 1; others=0, **Omnivore**=1; others=0) | **Wing morphology** (0=brachypterous; 1=dimorphic; 2=macropterous) | **Overwintering strategy** (imago hibernator=1; others=0) | Abiotic tolerance (0=hygro-; 1=meso-; 2=xerophilous) |
| Reptiles | **Body length** [cm] | **Diet breadth** (specialist=0, generalist=1) | **Movement distances** (reduced=0; moderate=1; high=2) | **Clutch size** (0=small; 1 = intermediate; 2=large) | Aquatic habitat affinity index (0=low;1=medium; 2=high) |

***Supplementary Table S4:*** Detailed description of **Amphibian** traits.

| ***Trait*** | ***Description and unit*** | ***Trait type*** | ***Sources*** |
|---|---|---|---|
| Body size | **Mean body length** (in cm) from the tip of the snout to the most posterior opening of the cloacal slit, snout–vent length (SVL). In the case of salamanders, total length measurements include body and tail. | Continuous | Baker et al. 2011<br>Beebee & Griffiths 2000<br>Frost 2021<br>Lips et al. 2003<br>Stevens et al. 2014<br>Trochet et al. 2014<br>amphibiaweb.org<br>animaldiversity.org<br>iucnredlist.org<br>research.amnh.org<br>Expert knowledge for single species scarcely documented. |
| Mobility | **Mobility.** Three categories: reduced (≤ 100 m), moderate (101 – 1000 m), and high (> 1000 m) levels of mobility in relation to regional pools. | Semi-continuous (0, 1, 2) | |
| Reproductive strategy | **Clutch size.** Three categories: small clutches (≤ 20 eggs), medium (21 – 300 eggs), and large (> 300 eggs). | Semi-continuous (0, 1, 2) | |
| Feeding | **Diet.** Two categories: specialists (those who ingest 1-2 food types), and generalists (consuming 3 or more food types). When this information is not available, use mouth size as a proxy of feeding traits, with larger mouths representing generalist species and smaller mouths representing specialists. | Semi-continuous (0, 1) | |
| Taxon specific | **Aquatic index.** Three categories: exclusively terrestrial, occupying ponds or multiple habitats, or exclusively riparian. | Semi-continuous (0, 1, 2) | |

***Supplementary Table S5:*** Detailed description of **Bat** traits.

| ***Trait*** | ***Description and unit*** | ***Trait type*** | ***Sources*** |
|---|---|---|---|
| Body size | **Forearm length** (in mm) | Continuous | Denzinger & Schnitzler 2013<br><br>Jung & Threlfall 2018<br><br>Expert knowledge for single species scarcely documented. |
| Mobility | Aspect ratio: the **ratio of wing span to wing area**. Higher aspect ratio enables fast, but less manoeuvrable flight. | Continuous | |
| Reproductive strategy | Bats were grouped into species specialized on certain **roosting requirements** (e.g., caves, foliage) or those that are flexible in their choice of roosting sites. | Categorical | |
| Feeding | Species were classified as those catching aerial insects in flight (**aerial hunters**) **and others**, which include gleaning prey from surfaces or the vegetation (gleaning), or perch hunting (the latter two categories were not abundant enough to keep separate and hence were merged for analysis). | Categorical | |
| Taxon specific | **Wing loading**: wing area per body mass<br><br>**Echolocation** (kHz): frequency of maximum amplitude or characteristic frequency (in the case of zero-cross-based recordings, i.e. Anabat recording systems) of echolocation calls.<br><br>**Habitat preference** classified as **foraging** in open habitats, or edge or cluttered habitats. The latter two were grouped due to insufficient numbers of species. The two categories were: foraging in open space=1; and others=0 (clutter, edge space). | Continuous<br><br>Continuous<br><br>Categorical (0,1) | |

***Supplementary Table S6:*** Detailed description of **Bee** traits.

| ***Trait*** | ***Description and unit*** | ***Trait type*** | ***Sources*** |
|---|---|---|---|
| Body size | Body size was given using the **inter-tegula distance**, ITD (in mm), given the two measures are highly correlated. ITD is the space between the two tegulae, which are the insertion points for each forewing. ITD measurements were obtained from the authors of each study, and are usually measured using an ocular micrometer or handheld calipers. | Continuous | Hinners et al. 2012<br>Normandin et al. 2017<br>Threlfall et al. 2015<br>Cariveau et al. 2016<br>Laurence Packer, York University, Toronto, Canada (pers comm)<br>Michael Batley, Australian Museum (pers comms)<br>Unpublished European bee trait database (compiler: Stuart Roberts; pollinator loss module of the EU- FP6 ALARM-project)<br>Expert knowledge for single species scarcely documented. |
| Mobility | **Inter-tegula distance**, ITD (mm) as above. | Continuous | |
| Reproductive strategy | **Sociality** was used as a proxy for reproductive strategy since it integrates several reproduction features (e.g., number of brood cells, gender organisation etc.). We classified sociality as ’solitary’ and ’other’, where the latter included eusocial, primitively-social or semi-social. | Categorical | |
| Feeding | **Tongue length,** categorised as short or long mouthparts. If species data were missing, tongue length was estimated using bee family and inter-tegula distance as per Cariveau et al. (2016), and subsequently assigned as short or long. | Categorical | |
| Taxon specific | Bees use a diversity of nesting locations or substrates, some of which can be heavily impacted upon by features of the urban environment. To simplify across the various **nesting strategies** that have been documented (Michener 2000) we classified species to the following:<br>Below ground (Below ground=1; others=0) Above ground (Above ground=1; others=0) Parasite (Parasite=1; others=0) | Categorical | |

***Supplementary Table S7:*** Detailed description of **Bird** traits.

| ***Trait*** | ***Description and unit*** | ***Trait type*** | ***Sources*** |
|---|---|---|---|
| Body size | Geometric mean of **body mass** average values for both sexes [in g]. | Continuous | Jetz et al. 2008<br>Sheard et al. 2020<br>Wilman et al. 2014 |
| Mobility | **Hand-wing index**, ratio of the difference between wing length (from carpal joint to tip of longest primary feather) and secondary length (from carpal join to tip of 1st secondary feather) by wing length [(wl-sl)/wl]. | Continuous | |
| Reproductive strategy | **Clutch size** [average number of laid eggs per nest]. | Continuous | |
| Feeding | Categorical **diet** assigned based on the dominant among five diet categories, based in the summed scores of individual diets [fruit-nectar (e.g., fruits, drupes, nectar, pollen, plant exudates, gums), invertebrates (e.g., shrimp, krill, crustaceans, molluscs, cephalopods, gastropods, insects, worms, etc.), plant-seed (e.g., seeds, nuts, grains, and other plant materials not included in fruit-nectar), vertebrates-scavenger (e.g., vertebrates, carrion, garbage, etc.), omnivorous (score of ≤ 50 of all specific categories)]. | Categorical | |
| Taxon specific | **Foraging strata index**. *Habitat* [0=aquatic; 1=terrestrial; 2=aerial] = (below surface + around surface) + 2*(ground + understory + mid high + canopy) + 3*(aerial);<br><br>*Aquatic*: [0 = does not forage in aquatic systems, 1 = forage on or just below water surface (<12.7cm), 2 = forage below water surfaces] = below surface + 2*around surface;<br><br>*Terrestrial* [0=does not feed in terrestrial systems, 1=feed on the ground, 2=feeds on the understory below 2 m, 3=feeds between 2 m and tree canopy, 4=feeds in the tree | Semi-continuous and categorical | |

| | |
|---|---|
| | canopy] = ground + 2* understory + 3 * mid high + 4 *canopy.<br>*Aerial* [0=does not feed well above vegetation or any structures, 1=feed well above vegetation or any structures]. |

***Supplementary Table S8:*** Detailed description of **Carabid beetle** traits.

| ***Trait*** | ***Description and unit*** | ***Trait type*** | ***Sources*** |
|---|---|---|---|
| Body size | Mean **body length** from the tip of the head to the tip of the abdomen (in mm) | continuous | Klaiber et al. 2017<br>Lindroth 1985, 1986<br>carabids.org<br>Expert knowledge for single species scarcely documented. |
| Feeding | Trophic guild. Three categories: *herbivore*, *carnivore*, ***omnivore***. | Categorical (0=no, 1=yes) | |
| Mobility | **Hind wing development**. Three categories: *brachypterous* (short-winged or wingless), *dimorphic* (short and long-winged individuals present in the same species), *macropterous* (long-winged). | Semi-continuous (0, 1, 2) | |
| Reproductive strategy | **Overwintering strategy**. Two categories: *spring breeder* (imago/adult hibernators, these species reproduce in the spring to early summer, their larvae develop in the summer and a new adult generation appears in the autumn, with these adults overwintering); *autumn breeder* (larval hibernators – these species reproduce in the summer or autumn and overwinter as larvae). | categorical (y/n) | |
| Taxon specific: Drought tolerance | **Tolerance to drought conditions**. Three categories: *hygrophilic* (wetness preference), *mesophlic* (intermediate preference) and *xerophilic* (drought preference). | Semi-continuous (0, 1, 2) | |

***Supplementary Table S9:*** Detailed description of **Reptile** traits.

| ***Trait*** | ***Description and unit*** | ***Trait type*** | ***Sources*** |
|---|---|---|---|
| Body size | Total **body length** for lizards, snakes, and crocodiles. **Carapace length** for turtles (in cm). | continuous | Stevens et al. 2014<br>reptile-database.org<br>animaldiversity.org<br>iucnredlist.org<br>research.amnh.org<br>Expert knowledge for single species scarcely documented. |
| Mobility | **Mobility.** Three categories: 0=reduced (≤ 100 m), 1=moderate (101 - 1000 m), and 2=high (> 1000 m) levels of mobility in relation to their year-round activities. | Semi-continuous (0, 1, 2) | |
| Reproductive strategy | **Clutch size.** Three categories: 0=small clutches (≤ 20 eggs), 1=medium (21 – 100 eggs), and 2=large (> 100 eggs). | Semi-continuous (0, 1, 2) | |
| Feeding | **Diet.** Two categories: specialists (those who ingest 1-2 food types), and generalists (consuming 3 or more food types). When this information is not available, use mouth size as a proxy of feeding traits, with larger mouths representing generalist species and smaller mouths representing specialists. | Semi-continuous (0, 1) | |
| Taxon specific: | **Aquatic index.** Three categories: 0=exclusively terrestrial, 1=occupying ponds or multiple habitats, or 3=exclusively riparian. | Semi-continuous (0, 1, 2) | |

***Supplementary Table S10***: Factor loadings on global climate PCA axes. Only the first four axes that were retained for further analyses are shown. PC1 = cold-warm temperature; PC2 = broad (e.g. deserts) – narrow diurnal range (e.g. tropics); PC3 = high-low variability of temperatures; PC4 = high-low seasonality of precipitation.

| | PC1 (55%) | PC2 (19%) | PC3 (9%) | PC4 (6%) |
|---|---|---|---|---|
| clim01: Annual Mean Temperature | **-0.284** | 0.197 | 0.049 | -0.063 |
| clim02: Mean Diurnal Range | -0.137 | **0.401** | 0.084 | 0.009 |
| clim03: Isothermality | -0.270 | 0.032 | -0.264 | -0.007 |
| clim04: Temperature Seasonality | 0.223 | 0.096 | **0.479** | 0.125 |
| clim05: Max Temperature of Warmest Month | -0.253 | 0.267 | 0.185 | -0.031 |
| clim06: Min Temperature of Coldest Month | **-0.296** | 0.121 | -0.089 | -0.094 |
| clim07: Temperature Annual Range | 0.174 | 0.216 | **0.512** | 0.141 |
| clim08: Mean Temperature of Wettest Quarter | -0.248 | 0.223 | 0.235 | 0.055 |
| clim09: Mean Temperature of Driest Quarter | -0.271 | 0.150 | -0.140 | -0.152 |
| clim10: Mean Temperature of Warmest Quarter | -0.259 | 0.249 | 0.180 | -0.037 |
| clim11: Mean Temperature of Coldest Quarter | **-0.294** | 0.141 | -0.084 | -0.082 |
| clim12: Annual Precipitation | -0.249 | -0.281 | 0.092 | 0.151 |
| clim13: Precipitation of Wettest Month | -0.245 | -0.193 | 0.014 | **0.370** |
| clim14: Precipitation of Driest Month | -0.163 | **-0.325** | 0.269 | -0.239 |
| clim15: Precipitation Seasonality | 0.030 | 0.150 | -0.200 | **0.677** |
| clim16: Precipitation of Wettest Quarter | -0.246 | -0.198 | 0.016 | **0.362** |
| clim17: Precipitation of Driest Quarter | -0.166 | **-0.327** | 0.265 | -0.234 |
| clim18: Precipitation of Warmest Quarter | -0.192 | -0.205 | **0.287** | 0.237 |
| clim19: Precipitation of Coldest Quarter | -0.185 | -0.266 | 0.001 | 0.015 |

***Supplementary Methods 1:*** Correlations among environmental variables. A = all taxa except birds; B = birds. Correlations between predictors are relatively low between urban land cover, forest land cover, latitude, and climate while being relatively high between percent cover and aggregation, as well as among different scales. Blue = positive correlations; Red = negative correlations.

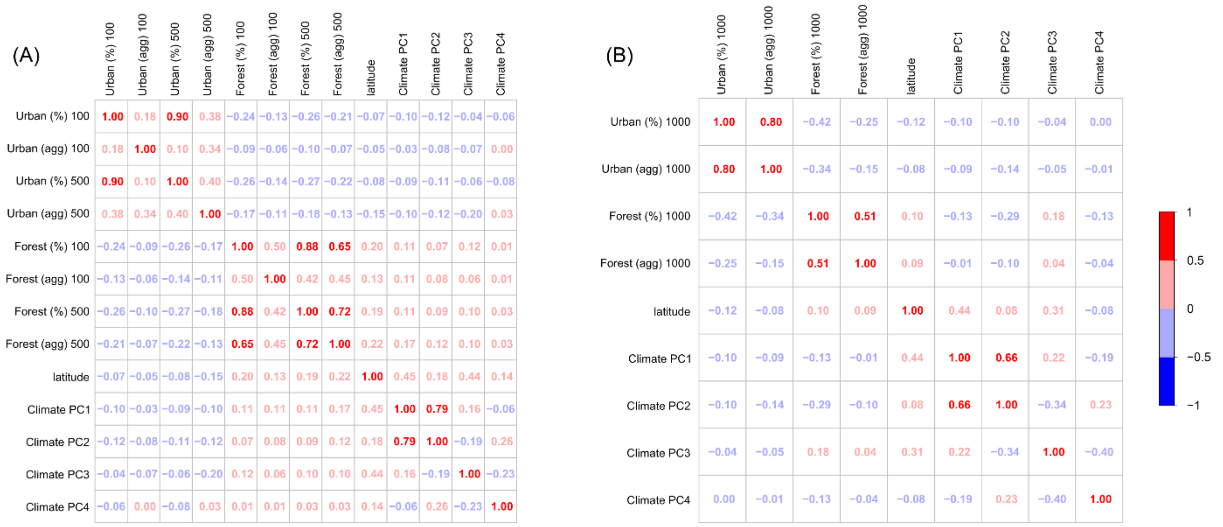
